# Dendritic Computation and the Fine Structure of Receptive Fields: A Model of V1 Neurons

**DOI:** 10.64898/2026.02.27.708525

**Authors:** Alessandro Paolo Bramanti

## Abstract

Dendrites are now recognized as nonlinear integrative compartments that can implement complex computations within single neurons, yet their specific contribution to cortical response diversity remains incompletely understood. In primary visual cortex (V1), neurons exhibit a broad range of response properties, including simple, complex, and end-stopped selectivity, but the mechanisms by which such diverse receptive field structures arise within a largely homogeneous population of pyramidal cells are still debated. This work presents a computational model in which these response classes emerge from dendritic integration of spatially organized excitatory and inhibitory inputs. In the model, synaptic inputs are distributed across dendritic branches according to their orientation preference and location within an underlying orientation map, and dendritic nonlinearities transform these inputs into somatic responses. By varying the relative distributions of excitatory and inhibitory inputs along dendrites, while keeping synaptic weights fixed, the model generates a continuum of response profiles ranging from phase-sensitive simple-like cells with segregated ON and OFF subfields to phase-invariant complex-like and end-stopped responses. The fine structure of model receptive fields, including the emergence of ON/OFF regions and end-stopping, arises from local differences in the balance and placement of excitatory and inhibitory inputs on dendritic branches. The model also predicts that modest changes in input organization can alter preferred orientation, consistent with reports of orientation plasticity in V1. These results provide a hypothesis-generating framework showing how dendritic input organization can constrain and shape V1 response properties within a single cell type, complementing classical feedforward and pooling-based accounts of these response classes.

## Introduction

Neurons in primary visual cortex (V1) exhibit a diverse range of response properties, including simple, complex, and end-stopped selectivity, which differ in their sensitivity to stimulus phase, spatial structure, and contrast boundaries^1–7^. Since the seminal work of Hubel and Wiesel, understanding how these response types arise from cortical circuitry has been a central goal of visual neuroscience. Modern classification work also indicates that simple and complex responses lie on a continuum rather than forming strictly separable categories^8^. Despite extensive experimental and theoretical work, the circuit mechanisms that give rise to this functional diversity within V1 remain an open question.

A broad class of models has been proposed to explain simple and complex cell properties, including feedforward convergence of lateral geniculate nucleus (LGN) inputs, nonlinear pooling mechanisms, and recurrent intracortical interactions^5–7,9–14^. Energy-model and normalization frameworks^5,6,11,14^ explain complex-like responses and contrast-invariant tuning through combinations of phase-shifted simple-like filters and divisive gain control, while recurrent network models account for contrast and contextual effects through intracortical amplification and inhibition^12,13^. These models successfully reproduce many aspects of V1 tuning and response invariances and have provided important insights into cortical computation. However, they typically treat cortical neurons as point-like integrators or rely on abstract nonlinearities at the level of the neuron as a whole. As a result, they place limited constraints on how synaptic inputs are organized across the dendritic arbors of individual neurons.

Pyramidal neurons in V1 possess elaborate dendritic trees, and a growing body of work indicates that dendritic integration is nonlinear and depends strongly on the spatial arrangement of excitatory and inhibitory synaptic inputs^15–21^. Dendritic spikes and other local nonlinear events can implement subunit-like computations that are sensitive to both the magnitude and location of synaptic input within the dendritic tree. This raises the possibility that dendritic organization itself plays an active role in shaping neuronal response properties, rather than merely serving as a passive substrate for synaptic summation. In particular, the relative placement of inputs along dendritic branches may constrain how visual features are integrated and transformed at the somatic level.

Experimental studies in V1 further suggest that the inputs to individual neurons are structured in ways that could interact with dendritic integration. Dendritic “spots” receiving inputs with different orientation preferences have been reported, and the distributions of excitatory and inhibitory inputs across orientation space and receptive field location differ between cells and across functional classes^22–27^. Moreover, the simple or complex character of V1 responses depends on the statistics of presynaptic input, and inhibitory inputs often exhibit broader or more heterogeneous tuning than excitatory inputs^24–26^. Together, these findings suggest that the spatial and functional organization of synaptic inputs, particularly the balance of excitation and inhibition along different dendritic branches, could be a key determinant of receptive field structure in V1. Throughout this work, the term dendritic topology refers to the relative spatial arrangement of excitatory and inhibitory synaptic inputs along dendritic branches (for example, proximal versus distal placement and local clustering), rather than to the position of neurons within the cortical orientation map.

The present work introduces a computational framework designed to explore this possibility. In particular, dendritic branches are treated as nonlinear computational subunits, and the spatial arrangement of excitatory and inhibitory synaptic inputs across these subunits is examined as a determinant of receptive field structure. The model consists of a single V1-like pyramidal neuron receiving orientation-selective inputs that are distributed across its dendritic arbor according to their tuning and position within an underlying orientation map. Dendritic integration is modeled with nonlinear conductances that capture essential features of local dendritic spikes, while abstracting from detailed biophysical mechanisms^15–19^. By varying the relative distributions of excitatory and inhibitory inputs along dendrites, the model generates a continuum of response profiles, including phase-sensitive simple-like responses with segregated ON and OFF subfields, phase-invariant complex-like responses, and end-stopped responses classically associated with “hypercomplex” cells. Importantly, these response types emerge within a single cellular architecture, without requiring qualitatively distinct neuron classes or an explicit hierarchy of simple and complex cells in the model.

This work is intended as a hypothesis-generating framework rather than a detailed reconstruction of V1 circuitry. The inputs are modeled phenomenologically, using orientation-selective filters as a simplified representation of the orientation structure present in afferent drive to V1^5,9,12,28–31^, and many aspects of cortical connectivity, including recurrent inputs, are deliberately abstracted. Within this simplified setting, the model demonstrates how dendritic input organization—specifically, the relative placement and balance of excitatory and inhibitory inputs along dendritic branches—can strongly constrain and shape V1-like response properties. The model is intended to provide a mechanistic account of classical receptive field properties—simple, complex, and end-stopped responses—within a single V1-like neuron, in a way that is directly testable with modern neurophysiological methods. The assumptions underlying the model, including the structure of presynaptic inputs and the form of dendritic nonlinearities, are summarized in a parameter table and discussed in relation to existing experimental evidence. The model yields a set of falsifiable predictions about the relationships between dendritic input organization and receptive field structure, and suggests that similar principles may operate across cortical areas where functional diversity arises within populations of morphologically similar pyramidal neurons.

## Methods

### Software

All simulations were performed using custom scripts written in MATLAB®^32^. The full simulation code and analysis routines are available in a public GitHub repository^33^, facilitating reproduction and modification of the model.

### Neuron model and equivalent dendritic circuit

Each model neuron is represented as a simplified pyramidal cell with five dendritic branches, each subdivided into four compartments. Dendritic integration is modeled using a phenomenological nonlinear conductance model inspired by previous work^18^. For computational efficiency, the total number of synaptic inputs per neuron was limited to 300. To obtain realistic somatic voltage amplitudes with this reduced number of inputs, conductance values were scaled while preserving their known dependence on distance from the soma. As a result, the model reproduces realistic somatic voltages but does not attempt to match absolute synaptic conductances or currents in detail.

The equivalent dendritic circuit follows the formulation in Jadi et al. (2012)^18^ and is described in full in the Supplementary Information. Parameters were adjusted so that the decay of peak depolarization and of spike threshold with distance along the dendrite matched the experimental measurements reported by Major et al. (2008) and Major et al. (2013)^16,19^. Somatic voltage was used as the main output variable, and a nominal firing threshold of -55 mV (15 mV above rest) was adopted for reference, although spiking dynamics were not explicitly modeled. For each neuron, the somatic membrane potential was rescaled by a constant factor such that the 80th percentile of its responses to the grating stimuli corresponded to 15 mV above rest (−55 mV), and this normalized voltage scale was used for all analyses and plots. Since spot stimuli typically evoke smaller somatic depolarizations than gratings, their receptive fields are interpreted in terms of relative voltage modulation across positions rather than absolute crossings of the nominal threshold. A schematic of the dendritic circuit and overall model architecture is provided in Figure 1.

**Figure 1.**
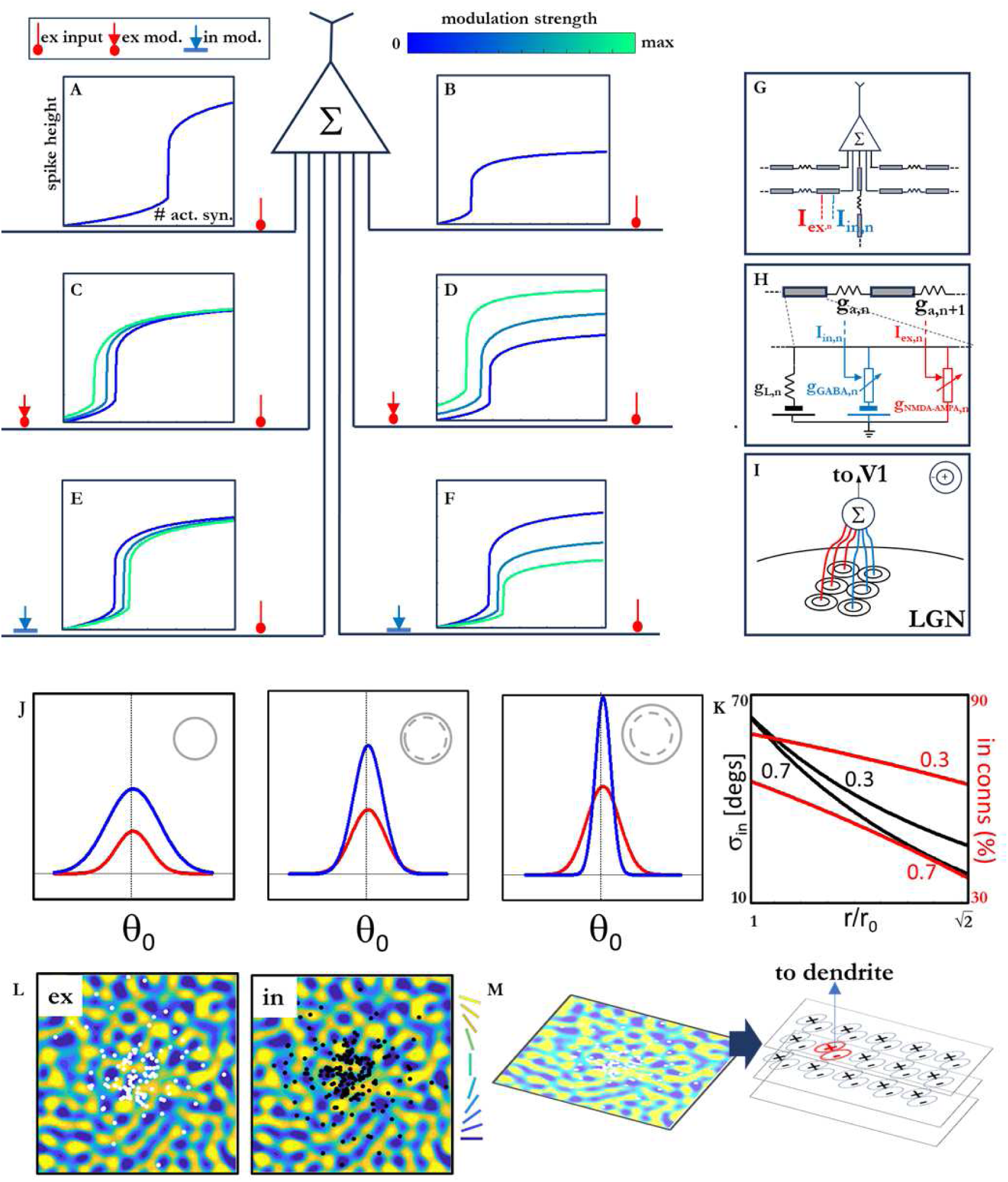
Graphical summary of the model. The effects of single or multiple inputs along the same dendritic branches are sketched in the top-left schematic (peak height measured at the soma vs input intensity curves, color code where appropriate indicates the modulation strength according to the scalebar). **A**, **B** Peak heights and thresholds for single inputs, respectively proximal and distal. **C**, **D** Effects of excitatory modulators, respectively distal and proximal. Both lower the threshold, the latter increase also the peak’s magnitude. **E**, **F** The same for inhibitory modulators, with specular effect with respect to the excitatory modulators. Realistic voltage and current scales for panels A to F are indicated in A. **G** 5-branch, 4-compartment dendritic structure of each neuron. **H**. Schematic of equivalent circuit for a single compartment. **I**. The elementary inputs are the output of Gabor filters capturing the local orientation in the image domain. **J**. Excitatory (red) and inhibitory (blue) Gaussian distributions of input orientations from Gabor filters, vs the dendritic radius (inset), centered around the preferred orientation θ_0_. In the insets, RF size (solid line) compared with minimum RF size (dashed line), drawing to scale. **K**. Inhibitory standard deviation and fraction of inhibitory connections vs the relative variation of the radius, for the two extreme values of the ex-to-in peak ratio (0.3 and 0.7, as marked in proximity of the curves). **L**. Excitatory and inhibitory connections (white and black dots, respectively) on the corresponding portion of the orientation map, for an example cell. **M**. When transforming from orientation to image space, both the spatial coordinates of the input and the local orientation detected (Gabor filter) must be determined.

### Circuit simulation

Because the excitatory conductance includes a nonlinear voltage dependence corresponding to NMDA-dominated supralinearity, the full dendritic circuit could not be solved analytically. Instead, for each input pattern the circuit was solved numerically by searching over a range of somatic voltages V_m_ and selecting the value that minimized the imbalance of currents at circuit nodes. Currents and local voltages in dendritic compartments were then computed using the analytical conductance expressions given in the Supplementary Information.

In this formulation, a single phenomenological excitatory conductance with sigmoidal dependence on membrane potential represents the combined effect of voltage-dependent NMDA channels and fast, approximately linear AMPA input, in line with previous modeling work^17–19^. Inhibitory input was modeled with a linear conductance. The model thus captures the essential nonlinearity of local dendritic spikes while abstracting from the detailed molecular contributions of different receptor types.

### LGN cells and Gabor filters

The feedforward drive to the model neurons represents the afferent input from LGN to V1. LGN cells were modeled as center–surround units, with a single-pixel central region and a surround of radius two pixels. Their response was normalized such that stimulation confined to the central pixel produced a response of 1, whereas uniform stimulation of the entire receptive field produced a response of 0.3.

To model the small-scale, orientation-selective inputs to V1 dendrites, the LGN responses were passed through a bank of linear Gabor filters. Each Gabor filter was elongated along its preferred orientation, with a radius of 6 pixels along the orientation axis and 4 pixels along the orthogonal axis, and consisted of two lobes (positive and negative). Filter outputs were linear and normalized. Twenty filters were used in total, corresponding to ten orientations from 0^∘^ to 162^∘^ in steps of 18^∘^, each with two polarities (positive and negative). Example filters are shown in Supplementary Figure SF-6.

These Gabor filters provide a phenomenological description of orientation-selective synaptic “spots” reported on V1 dendrites^22,23^, without specifying whether this selectivity arises from thalamic or intracortical sources. The model therefore treats the Gabor-filtered LGN outputs as the elementary inputs to dendritic synaptic sites.

### Orientation map generation

An orientation map was generated following the method of Rojer and Schwartz (1990)^34^. A two-dimensional white-noise image was band-pass filtered with a relative central spatial frequency r = 0.12 and relative bandwidth d = 0.06, both expressed relative to a maximum spatial frequency of 0.01. The resulting phase map was converted to orientation, and the angular values were quantized into ten discrete orientations from 0° to 162° in steps of 18°. Thus, each pixel in the map was assigned one of these orientation values.

Mapping between the orientation map and the image domain was assumed to be linear. A conversion factor between map space and image space was obtained by computing the average gradient of the orientation map, ∇_pw,av_, which quantifies the average rate of change of orientation per unit distance. The minimum dendritic radius in orientation-map coordinates was then chosen so that each dendritic compartment could, in principle, receive input from any of the discrete orientations. The corresponding minimum dendritic radius in image space was obtained using this conversion factor (see “Cell generation” below).

### Generation of excitatory and inhibitory orientation distributions

For each model neuron, the orientation preferences of excitatory and inhibitory inputs were drawn from Gaussian distributions centered on an assigned orientation 0_assigned_. The standard deviation of the excitatory distribution, σ_ex_, was randomly sampled from a uniform distribution between 40° and 50°, consistent with measurements in mouse V1^25,26^. The ratio between the peaks of the excitatory and inhibitory distributions was randomly chosen from a uniform distribution between 0.3 and 0.7, also based on Li et al. (2015)^26^.

The standard deviation of the inhibitory distribution, σ_in_, depended on dendritic radius. For the minimum cell radius r_O_, σ_in_ was at most 70^∘^. The dendritic radius for each cell was sampled from a uniform distribution between r_O_ and 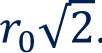, consistent with the limited range of dendritic tree sizes observed in V1 pyramidal neurons^35^. The number of excitatory connections at the minimum radius was determined, and this number was increased proportionally to dendritic area as the radius increased. The number of inhibitory connections was then set so that the total number of inputs per cell remained fixed at 300. For each cell, σ_in_ was adjusted to satisfy the chosen peak ratio and total number of inhibitory inputs given its radius. Mathematical details of this tuning procedure are provided in the Supplementary Information.

The ratio

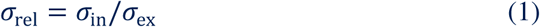

was used throughout the study as a summary parameter describing the relative breadth of inhibitory and excitatory orientation distributions. Parameter ranges and their sources (experimental data versus model assumptions) are summarized in Table 1.

**Table 1.** Model parameters and their sources.

| Parameter | Value | Animal model | Source refs. | Notes |
| --- | --- | --- | --- | --- |
| Dendritic input connections per cell | 300 | - | - | The constant number of input connections is an assumption of the model. The actual number (like the radius in pixels) has been chosen for good balance between computational speed and precision. |
| Minimum dendritic radius $r_0$ | 11 pixels | - | - | |
| Maximum dendritic radius | $r_0\sqrt{2}$ | Macaque | 35 | - |
| $\sigma_{ex}$ | $40^\circ \div 50^\circ$ | Mouse | 25, 26 | - |
| max $\sigma_{in}$ | $70^\circ$ | | | |
| ex-to-in peak ratio | $0.3 \div 0.7$ | Mouse | 26 | - |
| Number of ex connections grows proportionally to dendritic area | - | - | - | Assumption which, along with constant number of total connections, explains the variation of the inhibitory input distributions |
| Input distribution related to orientation map | - | - | - | Assumption, supported in particular by Refs. 22 (dendritic spots sensitive to orientation), 23 (composite structure of receptive fields), and 27 (relationship between orientation selectivity and position of neuron within orientation map). |

### Cell generation

Model neurons were generated in several steps:

1. *Cell position and assigned orientation.* A cell center was selected in both the orientation-map and image coordinate spaces, which are related by the conversion factor described above. The assigned orientation 0_assigned_ of the cell was taken as the orientation at the chosen position in the orientation map.
2. *Cell parameters and input distributions.* The dendritic radius of the cell was sampled between r_O_ and 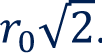. Given this radius, the standard deviation a_ex_ and the excitatory and inhibitory peak ratio were sampled within their allowed ranges. a_in_ was calculated, and the corresponding excitatory and inhibitory input distributions were determined as described in the previous section.
3. *Synaptic input generation in orientation–map space.* Excitatory and inhibitory synaptic inputs were generated in the orientation–map space according to the specified orientation distributions and a radial spatial distribution around the cell center. The radial distribution of synaptic sites followed an exponential decay with distance from the soma, such that synaptic density was highest near the center and decreased toward the periphery of the dendritic tree. Specifically, imagining a maximal dendritic radius of 240 µm, the decay length constant was set to 90 µm, so that the probability of placing a synapse decreased approximately as exp(−r/90) with radial distance r (details in the Supplementary Information).
4. *Mapping to image space and deduplication.* Synaptic input locations were mapped from orientation-map coordinates to image coordinates. Because of discretization and rounding, some inputs could map to the same pixel; duplicate connections were removed so that the effective total number of synapses per cell could be slightly less than the nominal value of 300.
5. *Dendritic tree and synapse assignment.* Five dendritic branches were created for each cell, with angular directions equally spaced by 72^∘^ starting from a randomly chosen initial angle. Each branch was subdivided into four equal segments (dendritic compartments). Synaptic inputs were assigned to dendritic branches and compartments based on proximity in image space. Each synapse retained the spatial coordinates and orientation preference of its corresponding Gabor filter.

All synaptic inputs had the same weight (normalized to 1), so that differences in response properties across neurons reflected differences in dendritic input organization rather than in synaptic strength.

### Receptive field mapping with spots

Receptive fields were mapped using bright and dark spots presented on a uniform background. For ON regions, a circular spot of radius 2 pixels and maximal luminance was moved across the receptive field on an otherwise dark background. For OFF regions, the polarity was reversed (dark spot on bright background). For each spot position, stimulus images were passed through the LGN model and Gabor filter bank as described above, and the resulting synaptic inputs were applied to the model neuron. The somatic membrane potential was recorded, and separate maps of responses to bright and dark spots were constructed.

### Grating stimuli and orientation tuning

Static sinusoidal grating stimuli with 100% contrast were used to measure orientation tuning. Gratings had spatial periods ranging from 6 to 50 pixels in steps of 2 pixels and phases from 0° to 340° in steps of 20°. As with spot stimuli, each grating was first processed by the LGN and Gabor filter bank, and the resulting synaptic drive was applied to the neuron model. For each orientation, the somatic responses to all spatial frequency and phase combinations were computed, and orientation tuning curves were obtained by taking, for each orientation, the average response across these combinations. This procedure was used consistently both for plotting the tuning curves shown in Figure 2B and for determining preferred orientations, defined as the orientations at which the averaged response was maximal.

**Figure 2.**
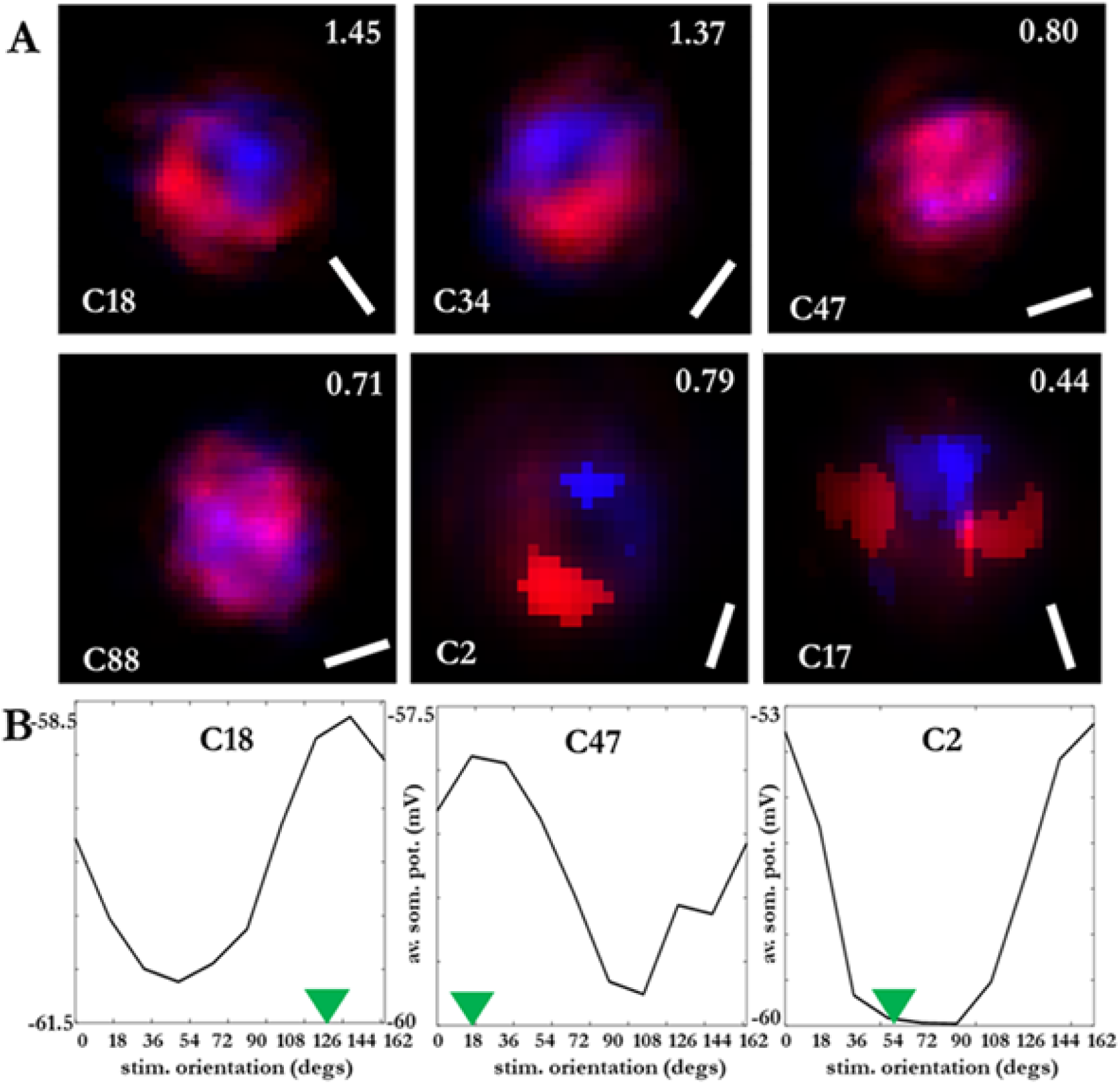
**A.** Spot maps of a selection of cells (ON and OFF regions in red and blue, respectively). Each map reports the cell index within the sample (lower-left corner), the nominal orientation (lower-right corner) and the value of σ*_rel_* (upper-right). C18 and C34 are simple, C47 and C88 are complex, C2 and C17 are recognized as hypercomplex cells after analysis. **B**. Mean somatic potential for three of the cells above elicited by a set of grating stimuli at variable orientations – the average is with respect to the periods and phases at each orientation. The green arrow indicates the nominal, distribution-based orientation, where the peak would be expected based on the distributions – the discrepancy in C2 is evident.

### Membrane orientation selectivity indices (mOSI_nom_ and mOSI_eff_)

Orientation selectivity was quantified using indices based on the somatic membrane potential. For each neuron, a membrane orientation selectivity index was defined as

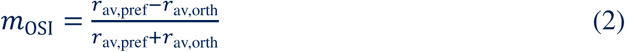

where r_av,pref_ is the average response at the preferred orientation and r_av,orth_ is the average response at the orientation orthogonal to the preferred one. Two indices were computed for each neuron. The nominal index, mOSI_nom_, used the assigned orientation 0_assigned_ (derived from the input distributions and orientation map) as the preferred orientation. The effective index, mOSI_eff_, used the orientation at which the grating-evoked response peaked as the preferred orientation. By construction, mOSI_eff_ is non-negative, whereas mOSI_nom_ can take negative values, indicating a preference for an orientation far from the assigned one.

### Effective receptive field diameter

An effective receptive field diameter was estimated from the spot-mapped responses and used to set bar width in end-stopping tests. For each neuron, the region in the spot map with responses greater than 1% of the peak response was approximated as circular, and the diameter of this circle was taken as the effective receptive field diameter.

### Bar stimuli and end-stopping tests

End-stopping was assessed using static bar stimuli presented on a uniform background. Bars had 100% contrast, with luminance polarity matched to the background (bright bars on a dark background and dark bars on a bright background), and were oriented at the same set of orientations used for grating stimuli (0° to 162° in 18° steps). For each neuron, bar width was set equal to the effective receptive field radius estimated from the spot-mapped responses (i.e. half of the effective receptive field diameter; see “Effective receptive field diameter”), so that the bar approximately matched the radius of the classical receptive field.

For a given neuron and bar orientation, bars were positioned so as to “enter” the receptive field from the border and extend across it. Specifically, for each orientation two opposite directions were probed independently: one bar started just outside the receptive field on one side, with its long axis aligned to the chosen orientation, and was elongated progressively across the field so that the length of its overlap with the receptive field increased from zero up to approximately the effective receptive field diameter; a second, specular bar performed the same sweep from the opposite side (for example, for horizontal bars, one entered from the left and elongated to the right, and the other entered from the right and elongated to the left). At each bar length and for each direction, the somatic membrane potential was computed and treated as a separate response curve.

Length-tuning curves were thus obtained, for each neuron, orientation, and direction of entry, by plotting the bar-evoked somatic depolarization as a function of bar length. Neurons were classified as end-stopped-like when at least one of these curves showed a pronounced increase in response with bar length, followed by a plateau over a limited range of lengths and a subsequent decline at longer lengths (see “Population-level response types”).

### Overlap index and inhibitory-to-excitatory indices

Spatial overlap between ON and OFF subregions was quantified using an Overlap Index (OI) based on the spot-mapped response fields. Let ON(x) and OFF(x) denote the somatic responses to bright (ON) and dark (OFF) spots, respectively, at position x in the receptive field map. The Overlap Index was defined as

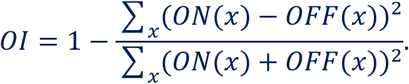

With this definition, O/ = 0 when the ON and OFF maps are completely segregated with no spatial overlap (for example, non-overlapping subfields of equal magnitude), and O/ = 1 when the ON and OFF maps are identical at every position. Intermediate values reflect partial overlap of ON and OFF subregions.

To assess the balance of inhibition and excitation in ON and OFF regions, an inhibitory-to-excitatory index was defined separately for bright and dark spots. For a given spot polarity s (bright or dark), the index

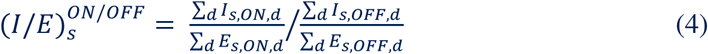

compares the inhibitory-to-excitatory ratio in ON versus OFF regions across dendritic branches, where d indexes the dendritic branches and /_s,ON,d_ and E_s,ON,d_ (respectively /_s,OFF,d_ and E_s,OFF,d_) are the total inhibitory and excitatory synaptic drives in the ON (respectively OFF) region on branch d for polarity s. In general, for bright spots the inhibitory-to-excitatory ratio is larger in the OFF than in the ON region, whereas for dark spots the opposite tends to hold. Consequently, 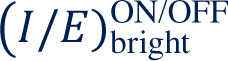 is typically less than 1 and 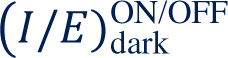 greater than 1.

### Parameter table

All model parameters, their values or ranges, and their origins (experimental measurement versus model assumption) are summarized in Table 1, together with species information and source references where applicable.

## Results

### Model input organization and population sampling

The model population consisted of 100 V1-like pyramidal neurons generated by sampling dendritic radii, excitatory and inhibitory orientation distributions, and synaptic placements according to the rules described in Methods. For each neuron, excitatory and inhibitory synaptic inputs were assigned orientation preferences and positions within an underlying orientation map and distributed across dendritic branches based on proximity. All synapses had equal weight in the baseline simulations, so differences in response properties arose solely from dendritic input organization and synapse placement. The receptive fields and responses of each neuron were characterized using spot mapping and static grating and bar stimuli. An overview of the model architecture, including the dendritic circuit, input generation, and orientation map, is shown in Figure 1./

### Example receptive fields along the σ_rel_ continuum

To illustrate how dendritic input organization shapes receptive field structure, a subset of neurons was examined in detail. Figure 2A shows spot-mapped receptive fields for representative cells spanning different values of the ratio σ_rel_ = σ_in_/σ_ex_, which summarizes the relative breadth of inhibitory and excitatory orientation distributions (Methods). For each cell, the bar in the corner indicates the assigned orientation derived from the input distributions and orientation map.

Cells with relatively large σ_rel_ exhibit receptive fields with clearly segregated ON and OFF subregions forming a band-like structure consistent with the classical definition of simple cells. In many of these neurons, the band orientation is approximately aligned with the assigned orientation and with the preferred orientation measured with grating stimuli, but the alignment is not exact and some cells show substantial deviations between these angles. Cells with σ_rel_ close to unity tend to show overlapping ON and OFF regions and phase-invariant responses more characteristic of complex cells. Cells with smaller σ_rel_ often display receptive fields with partially distinct ON and OFF regions whose orientation can differ substantially from the assigned orientation, and their responses to bars of varying length reveal pronounced end-stopping. Grating-based orientation tuning curves for example simple-like, complex-like, and end-stopped-like cells are shown in Figure 2B. For each orientation, these curves plot the average somatic response across spatial frequencies and phases, computed as described in Methods. For some cells with low σ_rel_, such as C2 and C17, the effective preferred orientation inferred from grating responses is nearly orthogonal to the orientation suggested by the input distribution, as reflected by negative values of mOSI_nom_ in Figure 3A. Across the population, simple-like cells showed nominal mOSI values between roughly -0.06 and 0.20 and effective values between about 0.006 and 0.20. Complex-like cells had nominal mOSI values from about -0.13 to 0.22 and effective values from about 0.02 to 0.23. End-stopped-like cells exhibited the broadest range, with nominal mOSI values from roughly -0.44 to 0.20 and effective values from about 0.04 up to 0.44. The corresponding bar-length responses for these two cells, illustrating their end-stopping behavior, are shown in Figure 3B. These examples motivate a population-level analysis of response types and their dependence on dendritic input organization.

**Figure 3.**
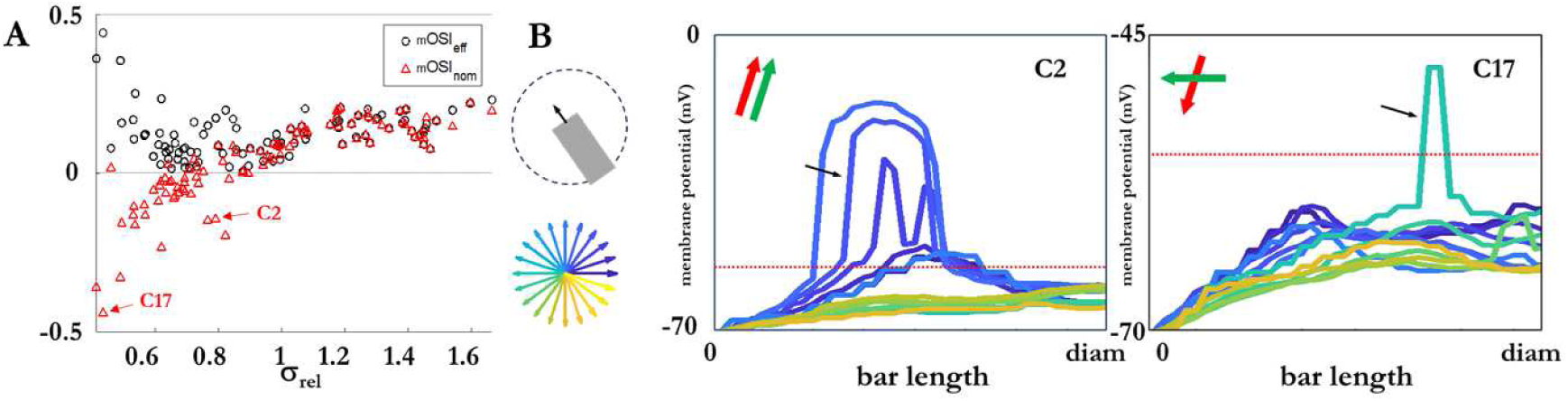
**A**. Membrane Orientation Sensitivity Indices (mOSI) both nominal and effective vs σ*_rel_*. C2 and C17, with negative *mOSI_nom_*, are evidenced. B. End-stopping tests in C2 and C17. The bar enters the RF along different angular orientations as indicated in the scheme and in the arrow diagram. In C2, some response is elicited for directions between 0 and 60° for bar lengths between roughly one quarter and one half of the cell’s diameter. C17 is more selective, reacting only to bars close to 180° and of precise length, around three quarters of the cell’s diameter. The black arrows indicate the curves which will be examined in detail later (Figure 6C). Insets: preferred cell orientation (red bar) and direction of highest maximum end-stopping effect (green arrow).

### Mechanisms of ON/OFF structure in simple-like and complex-like cells

To understand how dendritic input organization gives rise to simple-like and complex-like receptive fields, representative neurons from each class were analyzed in detail. In all cases, dendritic inputs are driven by the outputs of orientation-selective Gabor filters applied to bright and dark spots (Figure 4A), which appear as rings whose width and intensity differ between the two polarities.

**Figure 4.**
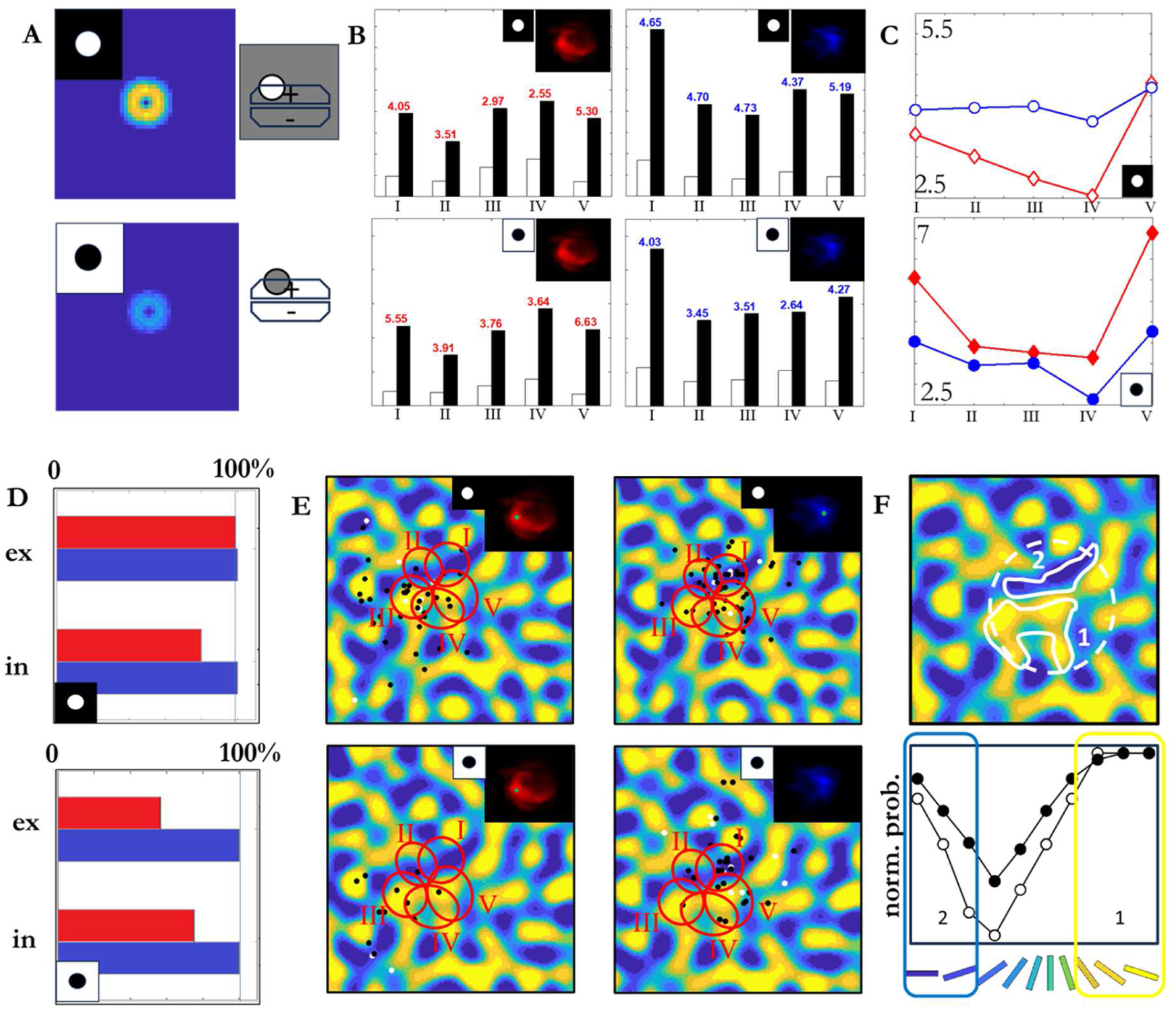
Analysis of simple cell C18. **A.** Gabor-filter ring-shaped responses to bright and dark spots, same color scale. The schemes on the right illustrate the different interactions between the spots and Gabor filters. **B.** Excitatory (white bars) and inhibitory (black bars) responses to both types of stimuli in both ON and OFF regions. The response is distinct by dendrite (Roman numerals) and limited to the most proximal compartment. Above each pair of bars, the inhibitory-to-excitatory ratio is reported. The same values are summarized in **C**. The curves with red diamonds (blue circles) are for ON (OFF) regions. Open (closed) symbols are for bright (dark) spots. **D.** Weighted averages of excited and inhibitory responses for the two types of spots in both regions (bars colored accordingly). **E.** Connections activated when each type of spot is in the point of maximum response of each region. The small green spot on the region marks the approximate position. Red ellipses: Domains of the most proximal compartments of each dendrite (numbered with Roman numerals). **F.** Local irregularity of the orientation map: regions 1 and 2 are the most abundant in connections bec/ause the corresponding orientations are well represented in the cell, as seen from the distributions (below) normalized at their peaks (white dots: excitatory, black dots: inhibitory). Yet, 2 is denser and appreciably smaller than 1.

Figure 4 illustrates the analysis for a simple-like cell (C18). The histograms in Figure 4B show, for each dendritic branch (most proximal compartment), the average excitatory and inhibitory input evoked by bright and dark spots in the ON and OFF regions of the receptive field. Above each pair of bars, the local inhibitory-to-excitatory ratio is indicated. In regions where the somatic response is suppressed, this ratio is consistently higher. For bright spots, the OFF region corresponds to locations where the inhibitory-to-excitatory ratio is high enough to prevent dendritic activation, whereas in the ON region the same spot elicits sufficient excitation at a lower inhibitory-to-excitatory ratio to generate a response. For dark spots, the situation is reversed. In the OFF region, the dark spot elicits enough dendritic activation because it tends to stimulate fewer connections and is therefore less likely to encounter large inhibitory-to-excitatory ratios. In the ON region, dark spots activate fewer excitatory connections than in the OFF region and do not recruit enough excitation to produce a significant response.

These branch-specific ratios are summarized in Figure 4C for both spot polarities and both regions. On average, the inhibitory-to-excitatory ratio is larger in the OFF than in the ON region for bright spots, and larger in the ON than in the OFF region for dark spots, but the exact values vary across branches due to local irregularities in the orientation map. Figure 4D shows the averages of excitatory and inhibitory inputs across branches, weighted by their contribution to the somatic potential, confirming that in C18 the bright spot induces an increase in inhibition and a decrease in excitation from ON to OFF regions, whereas the dark spot produces different trends but still yields distinct ON and OFF regions.

The spatial origin of these imbalances is illustrated in Figure 4E–F. For each polarity and region, the activated synaptic connections at the point of maximal response are plotted, with red ellipses marking the domains of the most proximal compartments (Figure 4E). In the ON region, the bright spot primarily activates excitatory inputs in specific dendrites at relatively low inhibitory-to-excitatory ratios, leading to a response. In the OFF region, the same spot activates predominantly inhibitory inputs in other dendrites, preventing a response. The dark spot shows complementary patterns.

These differences, in turn, depend on local irregularities of the orientation map, which, around C18, contains two prominent orientation subregions, labeled 1 and 2 (Figure 4F). Most of the cell’s proximal connections are drawn from these subregions, whose orientations coincide with or are close to the assigned orientation. Both subregions are rich in inhibitory connections, due to the large σ_rel_. However, subregion 1 is larger, so it features a lower spatial connection density. Subregion 2 is narrower and elongated, with a higher connection density, so that it is more likely to find many connections clustered in a single compartment with a high inhibitory-to-excitatory ratio. For these reasons, subregions 1 and 2 roughly coincide with the ON and OFF regions of the receptive field, respectively. A similar analysis for another simple-like cell (C34) is shown in Supplementary Figure SF-9 and yields analogous conclusions.

Complex-like cells behave differently (Figure 5). For the complex-like cell C47, the inhibitory-to-excitatory ratios in the most proximal compartments are substantially lower than in simple-like cells and show little dependence on spot polarity or receptive-field region (Figure 5A). The weighted averages of excitatory and inhibitory input in ON and OFF regions (Figure 5B) show no clear distinction between bright and dark spots or between the two regions, consistent with overlapping ON and OFF subfields and phase-invariant responses. This can be understood in terms of σ_rel_: for C47 and similar complex-like cells, inhibitory and excitatory inputs are more similarly distributed, making it less likely to find local regions where inhibition dominates sufficiently to suppress responses. Around the preferred orientation, inhibitory inputs are still more numerous than excitatory ones, but not enough to silence the neuron where excitation is strong. Supplementary Figure SF-10 shows analyses for two additional complex-like cells (C66 and C88), illustrating how both global input statistics and local orientation map structure interact with dendritic topology to determine whether a neuron expresses simple-like or complex-like receptive fields.

**Figure 5.**
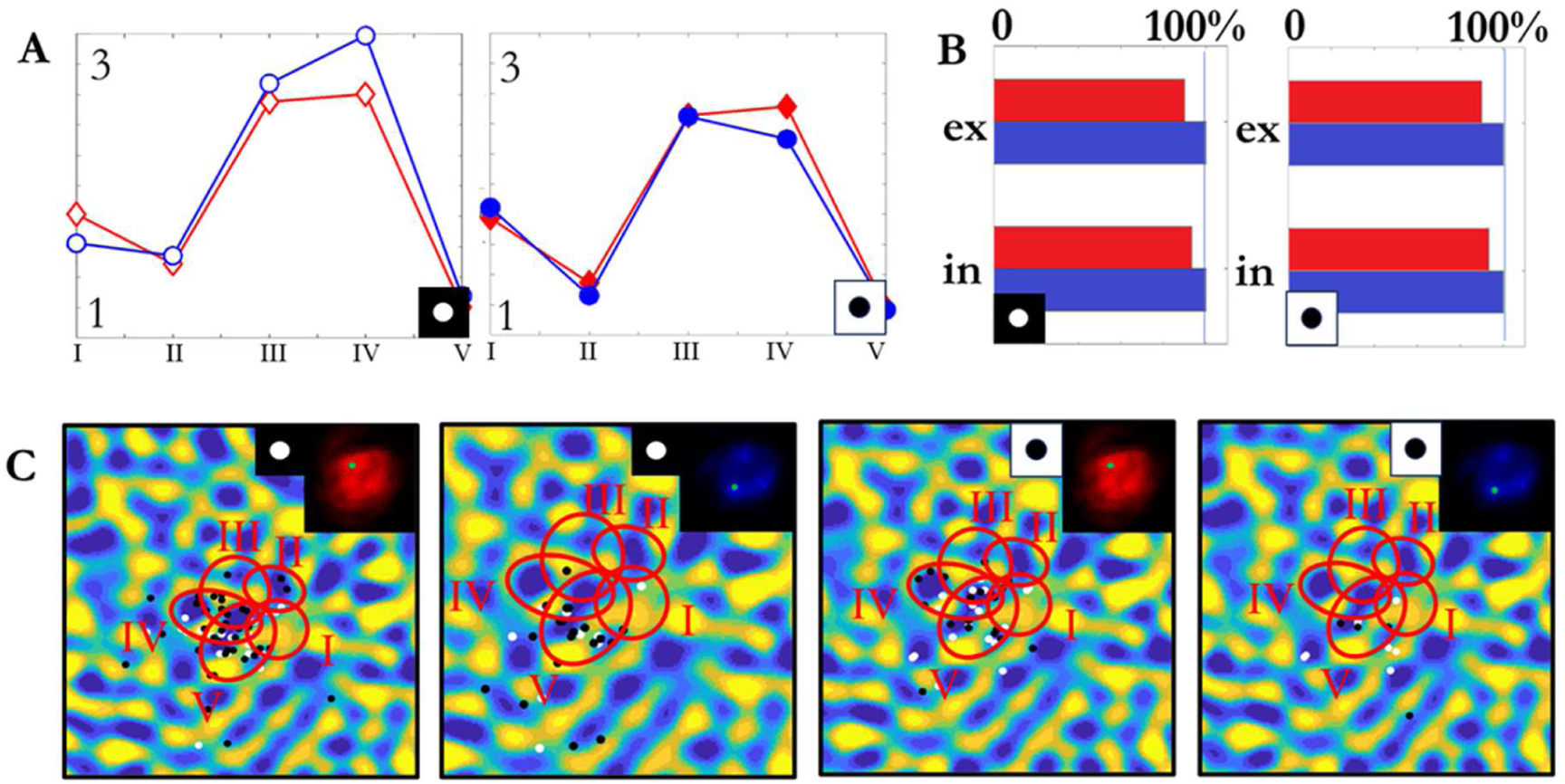
Analysis of complex cell C47. **A.** Inhibitory to excitatory ratios in the two regions with both stimuli, in the most proximal compartments of the five dendrites (same coding as in the previous figure). The values are remarkably lower than in simple cells and show negligible dependence on the stimulus. **B.** Weighted average of the excitatory and inhibitory inputs in the ON and OFF regions which, for a complex cell, overlap with each other (same coding as in the previous figure). There is no clear distinction between the responses to the two types of stimuli, nor between the two regions. **C**. Both stimuli were probed at the maximum response positions of the (mostly overlapping) ON and OFF regions. In all cases there is at least one compartment with considerable excitation vs inhibition, because the latter is more narrowly distributed.

### Mechanisms of orientation shifts and end-stopping in end-stopped-like cells

End-stopped-like cells in the model (e.g., C2 and C17) display two notable features: effective preferred orientations that can differ substantially from their assigned orientations, and strong end-stopping in bar-length responses. To investigate the mechanisms underlying these properties, the angular distributions of excitatory and inhibitory inputs across dendritic branches were examined (Figure 6A).

**Figure 6.**
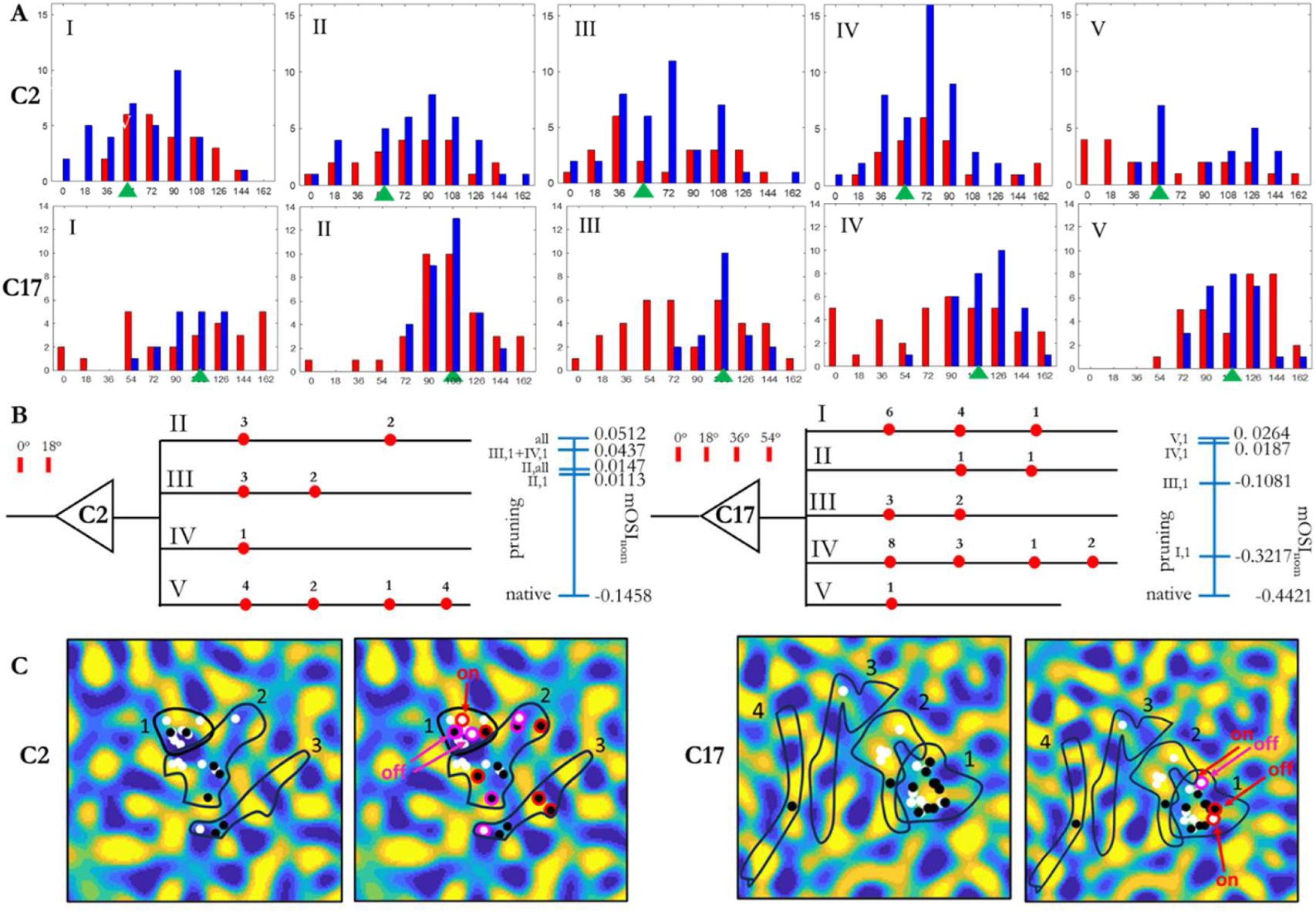
Analysis of end-stopped cells C2 and C17. **A** Angular distribution of excitatory (red) and inhibitory (blue) inputs in the different dendrites. **B** Diagrams showing the number of excitatory connections considered for pruning, distinct by dendrite. In C2 they are relative to 0 and 18 degrees (dendrite I is not represented for it does not contain any), in C17 to 0, 18, 36, and 54 degrees. On the right-hand side of each diagram the values of *mOSI_nom_* are reported vs the progressive pruning of excitatory connections. The dendrites and compartments (Roman and Arabic numerals, respectively) are pruned all at once for the considered orientations, as indicated on the left-hand side of the bar. The pruning is progressive (so previously pruned compartments are not restored). **C**. Two selected end-stopping-showing curves (one per cell, indicated by the black arrows in Figure 3) are analyzed vs the excitatory and inhibitory inputs in each of the dendritic branches. The curve is reported for each cell in the diagram above, the excitatory and inhibitory inputs are below for each branch (red and blue curves respectively). C2’s plateau is due to high excitatory-to-inhibitory rates in branches II and V, both showing some excitatory connections at small angles, unpaired by inhibitory connections. As for C17’s curve, the effect is to be attributed to branch I, where there is abundance of unpaired excitatory connections at small angles in the most proximal compartment.

Because these cells have low σ_rel_, the inhibitory orientation distribution is relatively narrow and the fraction of excitatory inputs is comparatively large. As a result, several dendritic branches receive excitation at orientations far from the assigned preferred orientation, with little or no inhibition at those orientations. Although the global input distributions are Gaussian, single branches can be strongly imbalanced due to local features of the orientation map. This produces distinct receptive-field subregions, as in simple-like cells, but here the imbalance is dominated by excitation rather than inhibition. Consequently, the nominal orientation derived from the global input distribution can differ markedly from the effective preferred orientation measured from responses, leading to negative values of mOSI_nom_ in some end-stopped-like cells. Similar, though typically smaller, mismatches between assigned and effective orientation also occur in some simple-like cells, indicating that orientation shifts in the model are a general consequence of dendritic input organization, with end-stopped-like cells representing an extreme case.

To assess how sensitive orientation preference is to changes in excitatory input, excitatory synapses at orientations far from the assigned value were selectively pruned, and the resulting shifts in mOSI_nom_ were measured (Figure 6B). For C2, excess excitatory connections at 0° and 18° were considered; for C17, excess excitation at 0°, 18°, 36°, and 54° was targeted. Pruning was performed at the level of entire compartments and was cumulative. In C2, eliminating excess excitation in the proximal compartment of dendrite V produced the largest single shift in orientation preference while removing only a small fraction of excitatory connections. C17 required more extensive pruning to achieve a comparable shift, consistent with its smaller σ_rel_ and broader distribution of excess excitation. Overall, these results show that the orientation preference of end-stopped-like cells can be altered substantially by relatively small changes in the distribution of excitatory inputs, suggesting a potential mechanism for orientation shifts observed after adaptation or learning.

End-stopping itself arises from the interaction between clustered, orientation-selective excitatory inputs on specific branches and the local recruitment of inhibition as stimuli extend beyond a critical spatial extent. Figure 6C examines, for each of C2 and C17, a single bar stimulus that elicits strong end-stopping. For C2, a bar at 36° elongated across the receptive field initially intercepts clusters of excitatory inputs on dendritic branch V that are largely unopposed by inhibition, producing a steep rise in somatic potential. Once the bar extends far enough to recruit additional inhibitory inputs in the same branch, the local dendritic potential drops and the somatic response falls abruptly, generating a plateau and subsequent decline. C17 shows a similar pattern, with end-stopping driven primarily by a single branch whose clustered excitatory inputs are selectively engaged by the bar. These analyses indicate that end-stopping in the model is a branch-specific phenomenon that depends on the precise spatial relationship between dendritic input clusters and the stimulus.

### Population-level response types

Each model neuron was classified as simple-like, complex-like, or end-stopped-like based on its receptive field structure and bar-length responses. Simple-like cells were defined as neurons with a clear Hubel & Wiesel–like band of alternating ON and OFF subregions in the spot map and no pronounced end-stopping, even when the band orientation deviated from the assigned orientation. Complex-like cells showed no clear band-like ON/OFF organization; although ON and OFF regions could be partially distinct, they did not form a consistent oriented subfield structure, and no strong end-stopping was observed. End-stopped-like cells exhibited a marked dependence on bar length at some orientation, characterized by a sharp rise in somatic potential, a plateau over a limited range of lengths, and a subsequent abrupt decline. These criteria are qualitative and reflect the fact that receptive field structure and length tuning vary continuously across the population; all cells were therefore inspected individually, and borderline cases are indicated in Supplementary Figure SF-7 together with their receptive fields and bar responses. Using these criteria, 25 neurons were classified as simple-like, 47 as complex-like, and 28 as end-stopped-like. Figure 7A summarizes the resulting distribution of functional types. Thus, the model does not produce a single canonical response type, but rather a continuum of receptive field structures spanning classical simple-, complex-, and end-stopped-like behavior within a single neuron architecture.

**Figure 7.**
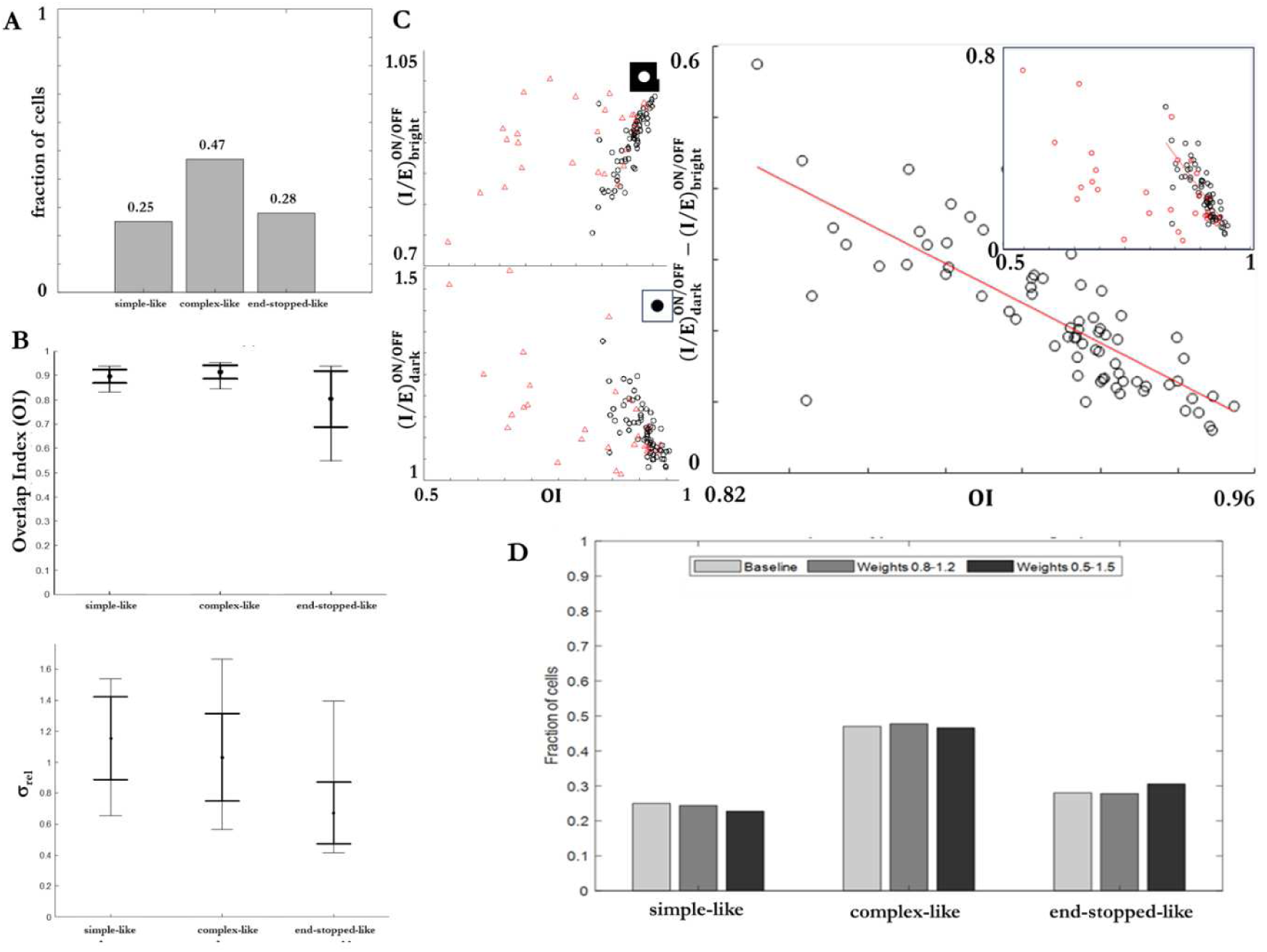
Population analysis and robustness. **A.** Fractions of simple-like, complex-like, and end-stopped-like cells in the model population (25, 47, and 28 cells, respectively). **B**. Summary of Overlap Index (OI, top) and σ_rel_ (bottom) across classes. For each class, mean values (black circles), ±SD (thick vertical whiskers) and min–max ranges (thin whiskers with horizontal caps) are shown. Simple-like and complex-like cells have relatively high OI, whereas end-stopped-like cells show lower OI on average. End-stopped-like cells also have smaller σ_rel_ than simple-like and complex-like cells, consistent with narrower inhibitory orientation distributions relative to excitation. **C**. Relationship between ON/OFF overlap and inhibitory–excitatory balance. Each point shows the Overlap Index (OI) and inhibitory-to-excitatory ratios 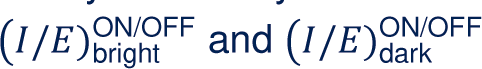 (Methods) for bright and dark spots in ON and OFF regions. In the scatter plots, simple-like and complex-like cells are shown as black circles, and end-stopped-like cells as red triangles. In the right-hand plot, the difference 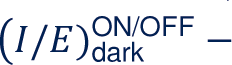 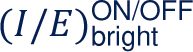 is plotted versus OI. The main panel displays only simple-like and complex-like cells (black circles), together with a linear regression fit (red line), highlighting a moderate approximately linear trend whereby neurons with more segregated ON and OFF subregions (lower OI) tend to show larger differences in inhibitory-to-excitatory balance between bright and dark spots. The inset shows all cells, including end-stopped-like cells (red triangles), which are more heterogeneous and do not follow this trend; some non-end-stopped cells also fall outside the main cluster, reflecting continuous variability in receptive field structure and dendritic input organization. **D**. Robustness of response-type distribution to synaptic weight perturbations. Fractions of simple-like, complex-like, and end-stopped-like cells are shown for the baseline condition (all synaptic weights equal to 1) and for populations in which each synaptic weight was multiplied by a random factor drawn from [0.8,1.2] or from [0.5,1.5]. Values for perturbed populations are averaged across five independent perturbation runs. The overall distribution of response types is largely preserved under modest weight perturbations, with only small shifts in class fractions, indicating that the emergence of simple-like, complex-like, and end-stopped-like responses does not depend on finely tuned synaptic weights.

The three classes differed in their average degree of ON/OFF overlap and in the relative breadth of inhibitory and excitatory orientation distributions (Figure 7B). Simple-like and complex-like cells both exhibited relatively high Overlap Index (OI) values, with mean OI of 0.896 ± 0.027 (mean ± SD; range 0.832–0.938) and 0.914 ± 0.027 (range 0.844–0.955), respectively, indicating substantial spatial overlap between their ON and OFF maps. End-stopped-like cells showed a lower mean OI of 0.804 ± 0.115 (range 0.548–0.939), consistent with more heterogeneous and often more segregated ON/OFF subfields in this group.

End-stopped-like cells also had the smallest mean σ_rel_ (0.672 ± 0.200, range 0.414–1.393), reflecting relatively narrow inhibitory orientation distributions compared to excitation. Simple-like cells had the largest mean σ_rel_ (1.155 ± 0.268, range 0.652–1.541), and complex-like cells occupied an intermediate range (1.031 ± 0.283, range 0.566–1.665). These trends indicate that smaller values of σ_rel_ favor the emergence of end-stopped-like responses, whereas larger values of σ_rel_ are associated with simple-like behavior and intermediate values with complex-like responses. At the same time, the OI and σ_rel_ distributions overlap substantially across classes, reflecting the influence of the detailed spatial arrangement of excitatory and inhibitory inputs along dendritic branches on receptive field structure and response type.

### Relationship between ON/OFF overlap and inhibitory–excitatory balance

The relationship between ON/OFF segregation and inhibitory–excitatory balance was examined across the population using the Overlap Index (OI) and inhibitory-to-excitatory indices 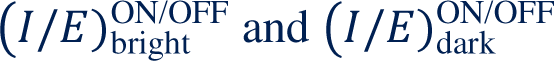 (Methods). Figure 7C shows the latter two indices as a function of OI. For bright spots on a dark background, 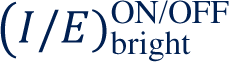 is generally less than 1, reflecting the fact that inhibition tends to outweigh excitation more strongly in OFF than in ON regions. For dark spots on a bright background, 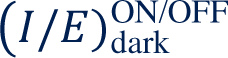 is typically greater than 1, consistent with higher inhibitory-to-excitatory ratios in ON than in OFF regions. Non-end-stopped cells (black points) tend to show a systematic relationship between these indices and OI, whereas end-stopped-like cells (red points) appear more heterogeneous. When the difference 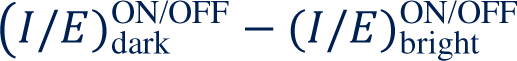 is plotted versus OI for simple-like and complex-like cells, a moderate approximately linear trend is observed: neurons with more segregated ON and OFF subregions (lower OI) tend to show larger differences in inhibitory-to-excitatory balance between bright and dark spots. End-stopped-like cells, by contrast, are more heterogeneous and do not follow this trend; many of them fall away from the main cluster of points, consistent with the additional constraints imposed by length tuning and branch-specific excitation. A small number of non-end-stopped cells also behave as outliers, reflecting continuous variability in receptive field structure and dendritic input organization. These cases are not analyzed separately here, but their receptive fields and response to bars are documented in Supplementary Figure SF-7. This pattern supports the idea that the degree of ON/OFF segregation is systematically related to how inhibition and excitation are distributed between ON and OFF regions, at least in simple-like and complex-like cells, whereas end-stopped-like cells reflect additional constraints imposed by length tuning and branch-specific excitation.

### Robustness to synaptic weight perturbations

To assess whether the distribution of response types depends sensitively on finely tuned synaptic weights, the same population of 100 neurons was simulated under random perturbations of synaptic strength. In each perturbation condition, every synaptic weight was multiplied by an independent random factor drawn either from [0.8,1.2] or from [0.5,1.5] with dendritic topology and input orientation distributions unchanged. For each condition, five independent realizations of the perturbed weights were simulated, and neurons were reclassified using the same criteria as in the baseline condition.

Under the milder perturbation ([0.8,1.2]), the fractions of simple-like, complex-like, and end-stopped-like cells changed only slightly across runs (Figure 7D). In each run, only a small number of neurons switched class, mostly between end-stopped-like and complex-like categories; one neuron switched between complex-like and simple-like in a single run. Two neurons that were essentially unresponsive to bar stimuli in the baseline condition became bar-responsive in all perturbed runs and were classified as complex-like. The effective preferred orientations of neurons shifted by approximately 4-6° on average (with standard deviations of about 9-15°), comparable to or smaller than the 18° sampling step used for orientation.

Under the larger perturbation ([0.5,1.5]), more classification changes were observed, but a substantial fraction of neurons retained their response type, and many changes again involved transitions between end-stopped-like and complex-like categories. Mean shifts in preferred orientation were on the order of 10-13°, with standard deviations of about 17-20°, still within one sampling bin. Taken together, these results indicate that the emergence of simple-like, complex-like, and end-stopped-like responses in the model does not depend on precisely tuned synaptic weights: dendritic input organization and the spatial arrangement of excitation and inhibition along branches are sufficient to generate a stable distribution of response types under modest weight variability.

## Discussion

### Summary and relation to previous models

This work examined how dendritic integration of spatially organized excitatory and inhibitory inputs can give rise to the diversity of response properties observed in primary visual cortex. In a single V1-like pyramidal neuron model, varying the relative orientation distributions and spatial placement of excitatory and inhibitory inputs along dendritic branches was sufficient to generate receptive fields and responses resembling classical simple-like, complex-like, and end-stopped-like cells. In this framework, simple-like, complex-like, and end-stopped-like responses, as traditionally defined in terms of ON/OFF structure and length tuning, emerge from differences in dendritic input organization within a common pyramidal neuron architecture. At the population level, the model produced a mixture of response types with different degrees of ON/OFF segregation, phase sensitivity, and end-stopping, even though all neurons shared the same basic cellular architecture and synaptic weights were held fixed.

Classical models of V1 emphasize feedforward convergence of LGN inputs, nonlinear pooling mechanisms, or recurrent intracortical interactions as the primary determinants of simple and complex cell properties^1,2,4–7,11–14^. Energy-model and normalization frameworks, for example, explain complex-like responses and contrast-invariant tuning through combinations of phase-shifted simple-like filters and divisive gain control^5,6,11,14^, while recurrent network models account for contrast and contextual effects through intracortical amplification and inhibition^12,13^. These frameworks successfully capture many aspects of orientation tuning, phase invariance, and contrast response, but they typically treat cortical neurons as point-like integrators or represent nonlinearities at the level of the neuron as a whole, without specifying how synaptic inputs are organized and processed across dendritic arbors.

The present model complements these approaches by explicitly incorporating dendritic topology—that is, the relative spatial arrangement of excitatory and inhibitory synaptic inputs along dendritic branches—as a computational variable. In this framework, simple-like, complex-like, and end-stopped-like responses need not reflect distinct neuron types or separate hierarchical stages; instead, they can emerge from differences in dendritic input organization within a common cellular design. The model therefore suggests that dendritic input topology, together with input statistics and intracortical circuitry, should be considered as an additional dimension in theories of V1 computation, alongside feedforward convergence, pooling, and normalization.

Other recent modeling work has incorporated dendritic nonlinearities and backpropagating action potentials into V1 models^36^, emphasizing network-level dynamics and a variational formulation that extends the standard linear–nonlinear framework. The present study instead focuses on a single multi-compartment pyramidal neuron and on how the spatial organization of excitatory and inhibitory inputs along its dendrites can generate a continuum of simple-like, complex-like, and end-stopped-like responses, together with the fine structure of receptive fields, within a common cellular architecture.

### Role of dendritic integration and input organization

Experimental work has established that dendritic integration in pyramidal neurons is highly nonlinear and depends on the spatial arrangement of synaptic inputs^15–21^. Dendritic spikes and other local nonlinear events can implement subunit-like computations, allowing dendritic branches to act as intermediate processing units between synapses and soma. In the present model, these nonlinearities interact with the structured distribution of excitatory and inhibitory inputs to shape receptive field structure and response diversity.

At the level of individual neurons, simple-like cells arise when inhibitory inputs are relatively broadly tuned and abundant (large σ_rel_), so that local clusters of inhibition can suppress responses in specific regions of the receptive field while sparing others, producing segregated ON and OFF subfields. Complex-like cells emerge when inhibitory and excitatory inputs are more similarly distributed (intermediate σ_rel_) and do not create strong local imbalances, reducing the impact of map irregularities and yielding responses that are more tolerant to spatial phase. End-stopped-like cells are associated with narrower inhibitory orientation distributions (small σ_rel_) and local excess excitation along particular dendritic branches, leading to strong responses within a limited spatial extent and suppression when stimuli extend beyond that range.

The population analyses show that these relationships are present at the group level but not deterministically tied to any single parameter: end-stopped-like cells, for instance, tend to have lower σ_rel_ and lower ON/OFF overlap than simple-like and complex-like cells, yet their distributions overlap substantially. This reflects the fact that global statistics such as σ_rel_ and OI capture only part of the story. The detailed dendritic topology—the specific placement of excitatory and inhibitory synapses along individual branches—further shapes receptive field structure and response type. In this sense, dendritic organization constrains and enriches the mapping from input statistics to functional properties, rather than merely implementing a fixed nonlinearity.

The model also suggests that orientation mismatches between assigned and effective preferred orientation can arise naturally from dendritic input organization. Even among simple-like cells, the orientation of the ON–OFF band is often only approximately aligned with the assigned orientation, and in some cases deviates substantially. End-stopped-like cells exhibit more pronounced and systematic mismatches, with effective preferred orientations that can approach orthogonality to the assigned value. Together with the pruning experiments, which show that small changes in the distribution of excitatory inputs can substantially shift orientation preference in end-stopped-like cells, these findings point to dendritic input structure as a potential substrate for orientation plasticity and map-level reorganization^27,37–40^.

### Limitations

The model makes several simplifying assumptions that should be considered when interpreting the results. First, it focuses on an isolated pyramidal neuron and does not include recurrent intracortical connections or feedback from higher visual areas, even though such inputs are abundant in V1 and are known to influence receptive field properties. In the present framework, these additional inputs could be incorporated as further sources of synaptic drive with their own spatial and orientation structure, but their contributions are not modeled explicitly here. As a consequence, phenomena that depend crucially on network dynamics, such as surround modulation or global normalization, are not addressed^14^.

Second, the afferent drive to the dendrites is modeled phenomenologically using Gabor–filtered LGN–like inputs that capture local orientation structure but do not distinguish between thalamic and intracortical sources. Experimental evidence for orientation–sensitive synaptic “spots” on V1 dendrites and for structured distributions of excitatory and inhibitory inputs across orientation and position^22–27^ justifies treating dendritic inputs as already organized in orientation space, but the precise mechanisms by which this organization arises remain unknown. The present model therefore abstracts this complexity into a simplified orientation map and associated input distributions^27,34^. More detailed models could incorporate specific LGN circuitry, intracortical pooling, or nonlinear pre–processing, but such extensions would primarily affect the amplitude and fine structure of dendritic inputs rather than the core dependence on their spatial arrangement.

Third, the dendritic integration mechanism is represented by a single phenomenological excitatory conductance with sigmoidal voltage dependence, intended to capture NMDA–dominated supralinearity together with fast AMPA input^16–19^. Inhibitory inputs are modeled with a linear conductance. This approach reproduces key qualitative features of distal and proximal inputs and dendritic spikes but does not resolve the separate contributions of different receptor types or voltage–gated channels. A more biophysically detailed model could test whether the qualitative relationships reported here are robust across different implementations of dendritic nonlinearity and conductance dynamics.

Finally, synaptic weights are held fixed in the main simulations, and variability in response properties arises from dendritic input organization and synapse placement. This choice isolates the effects of dendritic topology but does not address how synaptic weights and dendritic structure might be jointly shaped by development and plasticity. The robustness analysis indicates that modest random perturbations of synaptic strength do not qualitatively alter the distribution of response types, and even larger perturbations induce only moderate shifts in class membership and preferred orientation. However, the full interplay between synaptic weight tuning and dendritic input organization remains an important topic for future work.

The present model does not include explicit spike-generation dynamics, and all receptive field analyses are based on somatic membrane potential. Experimental work indicates that subthreshold membrane potential and spike-based measures of receptive field structure and orientation tuning are closely related, with spikes often reflecting a thresholded version of the underlying voltage response. Since the response regimes identified here are associated with large differences in somatic depolarization across stimuli and positions, they are expected to be preserved under reasonable spike-generation nonlinearities (for example, rectification above a nominal firing threshold).

### Falsifiable predictions

Although the model is conceptual and abstracts many aspects of V1 circuitry, it yields several specific, experimentally testable predictions:

#### 1. Dendritic input distributions and orientation maps

The model assumes that orientation-sensitive synaptic spots along dendrites are distributed in a manner related to the local orientation map. This predicts that, for neurons located near different regions of the orientation map (e.g., pinwheel centers versus orientation domains), the distribution of synaptic input orientations along dendrites should reflect the local map structure, rather than being uniform or random. This could be tested by combining in vivo functional imaging of dendritic inputs with local orientation map measurements^22–27^.

#### 2. Radius-dependent E/I balance

To account for the variation of inhibitory orientation width across cells, the model assumes that the fraction of inhibitory inputs decreases with dendritic radius while the central part of the inhibitory orientation distribution is preserved. This predicts that neurons with larger dendritic trees should, on average, receive a smaller proportion of inhibitory synapses than neurons with smaller trees, particularly in the tails of their orientation distributions, while maintaining similar excitatory tuning around the preferred orientation. Quantitative anatomical and functional mapping of E/I synapses as a function of dendritic arbor size^24–26,35^ could test this prediction.

#### 3. Dendritic E/I imbalance across branches

The model suggests that simple–like cells should exhibit local excess inhibition in some dendritic branches (contributing to suppressed regions in the receptive field), whereas end–stopped–like cells should exhibit local excess excitation in specific branches (contributing to strong, spatially limited responses). Complex–like cells are predicted to occupy an intermediate regime with more balanced dendritic trees. This could be tested by combining functional mapping of dendritic inputs with detailed reconstructions of excitatory and inhibitory synapses along individual branches, comparing branch–specific E/I ratios across cells classified as simple–like, complex–like, or end–stopped–like.

#### 4. Association of end-stopping with input organization

End-stopped-like behavior in the model is associated with relatively narrow inhibitory orientation distributions (low σ_rel_) and clustered excitatory inputs along specific branches. The model predicts that neurons exhibiting strong end-stopping should have dendritic input organization that reflects this pattern: narrower inhibitory tuning relative to excitation and pronounced clustering of similarly tuned excitatory inputs in the branches that drive end-stopping. High-resolution functional mapping of dendritic inputs in end-stopped neurons could test this association.

#### 5. Orientation plasticity and small input changes

The pruning experiments show that, in end-stopped-like cells, modest changes in the distribution of excitatory inputs (for example, weakening or strengthening a small fraction of synapses at certain orientations) can shift the effective preferred orientation substantially. This suggests that orientation preference in such cells could be particularly sensitive to changes in a relatively small subset of inputs, and that plastic changes in dendritic input organization could underlie orientation shifts observed after adaptation or learning, especially near pinwheel centers where input distributions are more heterogeneous^27,37–40^. Systematic measurements of orientation tuning before and after targeted dendritic plasticity would provide a direct test. These predictions are stated at the level of dendritic input organization and synaptic E/I balance and can be addressed using combinations of in vivo two-photon imaging, functional synaptic mapping, and anatomical reconstructions^20–27^.

### Broader implications for cortical computation

The present results highlight dendritic input organization as a powerful and biologically grounded mechanism for generating functional diversity in primary visual cortex. In this framework, neurons do not need to belong to distinct functional classes by design; instead, diverse response properties can emerge from variations in dendritic input organization within a common cellular architecture. This perspective aligns with growing experimental evidence that dendrites act as nonlinear integrative subunits^15–21^ and suggests that similar principles may operate across cortical areas, where differences in input organization rather than neuron type may underlie functional specialization. More generally, the findings support the view that cortical computation is not solely determined by synaptic convergence and output nonlinearities, but is also constrained by the spatial structure of synaptic inputs along dendrites.

### Implications for artificial and neuromorphic systems

Although the present work is focused on cortical computation, the results may also inform the design of artificial and neuromorphic systems. Most contemporary artificial neural networks implement computation primarily through learned synaptic weights, treating individual units as point-like integrators^28–31^. By contrast, the present model demonstrates that structured, nonlinear integration at the subunit (dendritic) level can strongly constrain functional selectivity, even when synaptic weights are not finely tuned. From this perspective, dendritic input organization can be viewed as a form of inductive bias that shapes computation through structure rather than parameter optimization alone. While translating such mechanisms into artificial systems remains an open challenge, incorporating spatially structured integration and subunit-like computation may offer a route toward more interpretable, robust, and biologically grounded architectures.

## Conclusions

This work presents a conceptual, hypothesis–generating model in which simple–like, complex–like, and end–stopped–like response properties emerge from dendritic integration of spatially organized excitatory and inhibitory inputs onto a single V1–like pyramidal neuron. By systematically varying dendritic input organization, particularly the relative breadth and placement of inhibitory and excitatory inputs along branches, the model reproduces a broad spectrum of receptive field structures and population behaviors using a common cellular architecture and fixed synaptic weights. These results support the idea that dendritic topology is a key determinant of cortical response diversity and provide a set of testable predictions about the relationship between dendritic input organization, synaptic E/I balance, and receptive field structure in V1 and beyond.

## Supporting information

Supplementary Methods and Figures. Full cell set data.

## References

1. Hubel, D.H., and Wiesel, T.N. (1959). Receptive fields of single neurons in the cat’s striate cortex. J. Physiol. 148, 574–591.

2. Hubel, D.H., and Wiesel, T.N. (1962). Receptive fields, binocular interaction and functional architecture in the cat’s visual cortex. J. Physiol. 160, 106–154.

3. DeAngelis, G.C., Ohzawa, I., and Freeman, R.D. (1993). Receptive-field structure in the cat’s primary visual cortex. I. Classical receptive fields. J. Neurophysiol. 69, 1091–1117.

4. Movshon, J.A., Thompson, I.D., and Tolhurst, D.J. (1978). Receptive field organization of complex cells in the cat’s striate cortex. J. Physiol. 283, 79–99.

5. Heeger, D.J., Simoncelli, E.P., and Movshon, J.A. (1996). Computational models of cortical visual processing. Proc. Natl. Acad. Sci. U.S.A. 93, 623–627.

6. Carandini, M., Heeger, D.J., and Movshon, J.A. (1997). Linearity and normalization in simple cells of the macaque primary visual cortex. J. Neurosci. 17, 8621–8644.

7. Rust, N.C., Schwartz, O., Movshon, J.A., and Simoncelli, E.P. (2005). Spatiotemporal elements of macaque V1 cells. Neuron 46, 945–956.

8. Mechler, F., and Ringach, D.L. (2002). On the classification of simple and complex cells. Vision Res. 42, 1017–1033.

9. Vintch, B., Movshon, J.A., and Simoncelli, E.P. (2015). A convolutional subunit model for neuronal responses in macaque V1. J. Neurosci. 35, 14829–14841.

10. Kouh, M., and Poggio, T. (2008). A canonical neural circuit for cortical nonlinear operations. Neural Comput. 20, 1427–1451.

11. Adelson, E.H., and Bergen, J.R. (1985). Spatiotemporal energy models for the perception of motion. J. Opt. Soc. Am. A 2, 284–299.

12. Troyer, T.W., Krukowski, A.E., Priebe, N.J., and Miller, K.D. (1998). Contrast-invariant orientation tuning in cat visual cortex: Thalamocortical input tuning and correlation-based intracortical connectivity. J. Neurosci. 18, 5908–5927.

13. Chance, F.S., Nelson, S.B., and Abbott, L.F. (1999). Complex cells as cortically amplified simple cells. Nat. Neurosci. 2, 277–282.

14. Carandini, M., and Heeger, D.J. (2012). Normalization as a canonical neural computation. Nat. Rev. Neurosci. 13, 51–62.

15. Poirazi, P., and Papoutsi, P. (2020). Illuminating dendritic function with computational models. Nat. Rev. Neurosci. 21, 303–321.

16. Major, G., Polsky, A., Denk, W., Schiller, J., and Tank, D.W. (2008). Spatiotemporally graded NMDA spike/plateau potentials in basal dendrites of neocortical pyramidal neurons. J. Neurophysiol. 99, 2584–2601.

17. Behabadi, B.F., Polsky, A., Jadi, M., Schiller, J., and Mel, B.W. (2012). Location-dependent excitatory synaptic interactions in pyramidal neuron dendrites. PLoS Comput. Biol. 8, e1002599.

18. Jadi, M., Polsky, A., Schiller, J., and Mel, B.W. (2012). Location-dependent effects of inhibition on local spiking in pyramidal neuron dendrites. PLoS Comput. Biol. 8, e1002550.

19. Major, G., Larkum, M.E., and Schiller, J. (2013). Active properties of neocortical pyramidal neuron dendrites. Annu. Rev. Neurosci. 36, 1–24.

20. Smith, S.L., Smith, I.T., Branco, T., and Häusser, M. (2013). Dendritic spikes enhance stimulus selectivity in cortical neurons in vivo. Nature 503, 115–120.

21. Lavzin, M., Rapoport, S., Polsky, A., Garion, L., and Schiller, J. (2012). Nonlinear dendritic processing determines angular tuning of barrel cortex neurons in vivo. Nature 490, 397–401.

22. Jia, H., Rochefort, N.L., Chen, X., and Konnerth, A. (2010). Dendritic organization of sensory input to cortical neurons in vivo. Nature 464, 1307–1312.

23. Fournier, J., Monier, C., Levy, M., Marre, O., Sári, K., Kisvárday, Z.F., and Frégnac, Y. (2014). Hidden complexity of synaptic receptive fields in cat V1. J. Neurosci. 34, 5515–5528.

24. Mariño, J., Schummers, J., Lyon, D.C., Schwabe, L., Beck, O., Wiesing, P., Obermayer, K., and Sur, M. (2005). Invariant computations in local cortical networks with balanced excitation and inhibition. Nat. Neurosci. 8, 194–201.

25. Liu, B., Li, Y., Ma, W., Pan, C., Zhang, L.I., and Tao, H.W. (2011). Broad inhibition sharpens orientation selectivity by expanding input dynamic range in mouse simple cells. Neuron 71, 542–554.

26. Li, Y., Liu, B., Chou, X., Zhang, L.I., and Tao, H.W. (2015). Synaptic basis for differential orientation selectivity between complex and simple cells in mouse visual cortex. J. Neurosci. 35, 11081–11093.

27. Nauhaus, I., Benucci, A., Carandini, M., and Ringach, D.L. (2008). Neuronal selectivity and local map structure in visual cortex. Neuron 57, 673–679.

28. Kriegeskorte, N. (2015). Deep neural networks: A new framework for modeling biological vision and brain information processing. Annu. Rev. Vis. Sci. 1, 417–446.

29. Kriegeskorte, N., and Diedrichsen, J. (2016). Inferring brain-computational mechanisms with models of activity measurements. Philos. Trans. R. Soc. Lond. B Biol. Sci. 371, 20160278.

30. Riesenhuber, M., and Poggio, T. (1999). Hierarchical models of object recognition in cortex. Nat. Neurosci. 2, 1019–1025.

31. Serre, T., Wolf, L., Bileschi, S., Riesenhuber, M., and Poggio, T. (2007). Robust object recognition with cortex-like mechanisms. IEEE Trans. Pattern Anal. Mach. Intell. 29, 411–426.

32. MATLAB (2022). Version 9.13.0.2049777 (R2022b). Natick, Massachusetts: The MathWorks Inc.

33. Bramanti, A.P. (2025). V1 cells – Generation and Characterization. GitHub repository, https://github.com/alessandro-bramanti/V1-cells---Generation-and-Characterization.

34. Rojer, A.S., and Schwartz, E.L. (1990). Cat and monkey cortical columnar patterns modeled by bandpass-filtered 2D white noise. Biol. Cybern. 62, 381–391.

35. Oga, T., Okamoto, T., and Fujita, I. (2016). Basal dendrites of layer-III pyramidal neurons do not scale with changes in cortical magnification factor in macaque primary visual cortex. Front. Neural Circuits 10, 74.

36. Rentzeperis, I., Prandi, D., and Bertalmio, M. (2025). A neural model for V1 that incorporates dendritic nonlinearities and backpropagating action potentials. J. Neurosci. 45(43), e1975242025.

37. Dragoi, V., Shapley, R.M., and Sur, M. (2000). Adaptation-induced plasticity of orientation tuning in adult visual cortex. Nature 403, 272–276.

38. Yao, H., and Dan, Y. (2001). Stimulus timing-dependent plasticity in cortical processing of orientation. Neuron 32, 315–323.

39. Dragoi, V., Sharma, J., Miller, E.K., and Sur, M. (2002). Dynamics of neuronal sensitivity in visual cortex and local feature discrimination. Nat. Neurosci. 5, 883–891.

40. Dragoi, V., Rivadulla, C., and Sur, M. (2001). Foci of orientation plasticity in visual cortex. Nature 411, 80–86.

