## Supplementary Methods and Figures. Full cell set data. for "Dendritic Computation and the Fine Structure of Receptive Fields: A Model of V1 Neurons"

### Supplementary Information

#### *SI-1 Parameters in the equivalent circuit.*

The equivalent circuit of the dendrite is as follows:

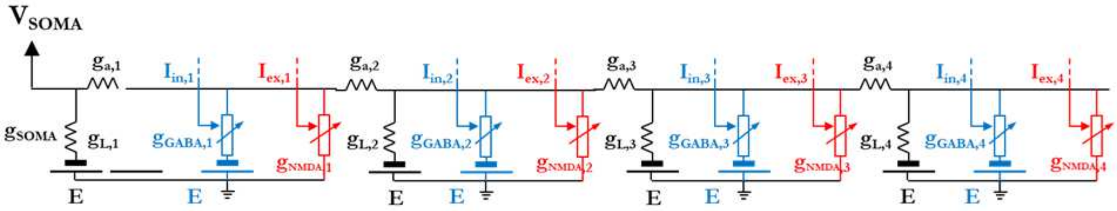


**Figure SF-1** – Equivalent circuit of 4-compartment dendrite, including the somatic conductance at which the voltage is measured.

Dendritic spiking is dominated by NMDA conductance[16,18,19]. Accordingly, a constant NMDA/AMPA ratio fits the experimental data well, but the nonlinearity of dendritic spikes originates primarily from the former[21]. Thus, each cell includes in each dendritic compartment a single excitatory, NMDA-type conductance[16,17].

$$g_{NMDA,n} = \frac{\gamma_{NMDA,n} I_{ex,n}}{1+\exp\left(-\frac{V_m+23.5}{12.5}\right)} \qquad \text{(SE1)}$$

in which the index *n=1,2,3,4*, as in the other equations, indicates the compartment number, $\gamma_{NMDA,n}$ is the compartmental conductance per unit synaptic current, and $I_{ex,n}$ is the sum of the excitatory currents (outputs of Gabor filters) afferent to the *n*-th compartment. The numerical constants are as in Ref. 16 (Figure S4, Supporting Information) and Ref. 18 (Figure S3, Supporting Information). The time-dependent part of the NMDA conductance is here neglected.

[1] STMicroelectronics srl, c/o Campus Ecotekne, via Monteroni 165, I-73100 Lecce, Italy,

In addition, each compartment includes a GABAergic conductance

$$g_{GABA,n} = \gamma_{GABA,n} I_{in,n} \qquad \text{(SE2)}$$

depending on the sum of afferent inhibitory currents. Both currents, $I_{ex,n}$ and $I_{in,n}$, summarize the effects of all (respectively excitatory and inhibitory) synapses activated in compartment *n*. In this model, they are the combined outputs of Gabor filters drawing on the LGN input and are, then, expressed in arbitrary input units (*a.i.u.*). As already mentioned, $\gamma_{NMDA,n}$ and $\gamma_{GABA,n}$ are then conductances per *a.i.u.*.

All the batteries supply

$$E = -70\ mV \qquad \text{(SE3)}$$

while no battery (zero voltage) is in series with the NMDA conductance, as in Jadi et al. (2012), Ref. 18.

*Calculation of the threshold excitatory current.*

The threshold of the dendritic spike at each compartment is determined in the absence of inhibition and with all NMDA conductances inactive except the one in the compartment itself, so that the equivalent circuit becomes:

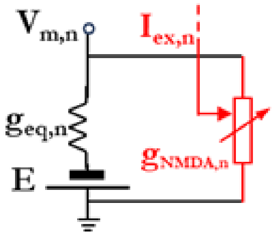


**Figure SF-2** – Equivalent circuit of one compartment: $g_{eq,n}$ is the equivalent conductance in parallel to the n-th NMDA conductance, controlled by the n-th excitatory current, while $V_{m,n}$ is the membrane voltage.

in which $g_{eq,n}$ is the equivalent conductance of the rest of the dendritic branch, including both the distal and proximal part – constant because the rest of the circuit, in conditions of zero excitation and inhibition, becomes linear.

According to Ref. 16, somatic voltage peaks range approximately from 23 to 3 mV, moving the origin location from 60 to 240 μm, with an 87 μm length constant. Considering four compartments, 60-μm spaced apart from each other and covering that distance range, the somatic peak originated in the n-th compartment is well approximated by:

$$V_{peak,n} = 23 \exp\left(-\frac{60(n-1)}{87}\right). \tag{SE4}$$

The compartmental NMDA conductance decreases remarkably from soma to periphery. According to Ref. 16, the variations in peaks and threshold vs distance are to be attributed substantially to NMDA conductance, so that we can assume $\gamma_{NMDA}$ to decay according to the same 87-μm length constant. With a proximal conductance as large[16] as 173 nS $\gamma_{NMDA}$, expressed in nS per unit current, can be written as:

$$\gamma_{NMDA,n} = 173 \exp\left(-\frac{60(n-1)}{87}\right). \tag{SE5}$$

The inhibitory conductances share the same exponential profile:

$$\gamma_{GABA,n} = \frac{\gamma_{NMDA,n}}{2}. \tag{SE6}$$

The highest somatic voltages obtainable without eliciting a dendritic spike range from 9.1 to 1.3 mV, proximal to distal with a decay length of 96 μm[16]:

$$V_{subth-soma,n} = 9.1 \exp\left(-\frac{60(n-1)}{96}\right). \tag{SE7}$$

The subthreshold voltage at the compartment's site can be estimated from (SE4), assuming for simplicity that the dendritic peaks at generation are all as large as 70 mV above rest potential and that the subthreshold voltage undergoes the same attenuation as the peak:

$$V_{subth-comp,n} = 70x9.1 \exp\left(-\frac{60(n-1)}{96} + \frac{60(n-1)}{87}\right). \tag{SE8}$$

The actual amplitude can be lower than that, so that making this assumption eventually leads to dendritic peaks at the soma slightly lower than intended. A correction would be possible through repeated estimation but would not bring any substantial improvement as long as the dependence on the distance to soma is correct, because the somatic voltage should be anyway corrected as indicated below.

The iontophoretic current needed to elicit a spike is as large as 200 nA proximally, decaying 4.9-fold to the distal compartments. Assuming the compartmental currents are at threshold are proportional, though smaller, the current profile expressed in nA can be approximated by:

$$I_{th,n} = 2\exp\left(-\frac{60(n-1)}{120}\right), \qquad \text{(SE9)}$$

assuming that, when the first compartment is at threshold, the summed transmembrane current is as large as 2 nA.

The equivalent conductance seen by the n-th compartment is, then:

$$g_{eq,n} = \frac{I_{th,n}}{V_{subth-comp,n}}, \qquad \text{(SE10)}$$

while the input excitatory current at threshold, through (SE1) is:

$$I_{ex-th,n} = \frac{I_{th,n}}{V_{subth-comp,n}-E_m}\left[1+\exp\left(-\frac{V_{subth-comp,n}-E_m+23.7}{12.5}\right)\right]. \qquad \text{(SE11)}$$

*Calculation of the somatic attenuation.*

To tune the somatic amplitude of the dendritic peaks vs the position, the n-th cell of Figure SF-2 can be expanded as follows:

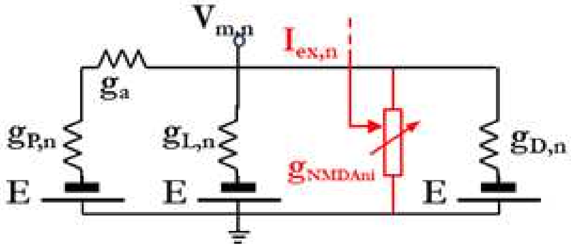

**Figure SF-3** – Expanded equivalent circuit of a compartment: impedance $g_{eq,n}$ has been exploded into its dependence on the n-th leakage and axial conductances as well as the n-th equivalent conductances of the proximal and distal parts of the dendritic branch, $g_{P,n}$ and $g_{D,n}$.

where $g_{P,n}$ and $g_{D,n}$ are the equivalent proximal and distal conductances respectively, i.e. those summarizing the effects of the other compartments, respectively towards and far from the soma, and $g_a$ is the axial conductance connecting the compartment to its more proximal neighbor – or, the most proximal compartment to the soma.

The conductances can be calculated for all the cells, starting with the N-th (most distal compartment) and moving backward to the second (n=2), observing that:

$$g_{D,N} = 0; \quad g_{D,n} = \frac{(g_{D,n+1}+g_{L,n})\cdot g_{a,n}}{g_{D,n+1}+g_{L,n}+g_{a,n}} \text{ for n<N,} \qquad \text{(SE12)}$$

$$\frac{g_{a,n}\cdot g_{P,n}}{g_{a,n}+g_{P,n}} + g_{L,n} + g_{D.n} = g_{eq,n}, \qquad \text{(SE13)}$$

$$\frac{g_{a,n}}{g_{a,n}+g_{P,n}} = \frac{p_n}{p_{n-1}}, \qquad \text{(SE14)}$$

$$g_{P,n} = g_{eq,n-1} - \frac{g_{a,n}(g_{L,n}+g_{D,n})}{g_{a,n}+g_{L,n}+g_{D,n}}. \qquad \text{(SE15)}$$

Equation (SE14) states that each cell introduces an attenuation of the voltage peak according to the term in the left-hand member. The product of all the attenuation terms from the *N*-th cell to the soma must equal the desired attenuation according to the (SE4). The system is solved by:

$$g_{a,n} = \frac{g_{eq,n}g_{eq,n-1}p_n}{n-g_{eq,n-1}p_n^2}, \qquad \text{(SE16)}$$

$$g_{P,n} = \frac{-g_{eq,n} g_{eq,n-1}(p_n - 1)}{g_{eq,n} - g_{eq,n-1} p_n^2}, \quad \text{(SE17)}$$

$$g_{L,n} = \frac{g_{eq,n}^2 - g_{eq,n}(g_{eq,n-1} p_n + g_{D,n}) + g_{eq,n-1} p_n^2 g_{D,n}}{g_{eq,n} - g_{eq,n-1} p_n^2}. \quad \text{(SE18)}$$

When n=1, the right-hand member of Equation (SE13) is simply the attenuation of the most proximal cell – in this case such that the plateau is proportional to the 23 mV reported in the literature. However, Equation (SE13) cannot be written in this form because $g_{P,1}$, the conductance at the soma, cannot be expressed as the ratio to the conductance of any other more proximal cell. The choice is to set:

$$g_{L,1} = g_{eq,1} \cdot mean(g_{L,n} / g_{eq,n})_{n>1}. \quad \text{(SE19)}$$

Then:

$$g_{S,1} = \frac{(g_{eq,1} - g_{L,1} - g_{D,1})}{p_1} \quad \text{(SE20)}$$

/

$$g_{a,1} = \frac{(g_{eq,1} - g_{L,1} - g_{D,1}) \cdot g_{P,1}}{g_{P,1} - (g_{eq,1} - g_{L,1} - g_{D,1})} \quad \text{(SE21)}$$

*Parameter values*

The values chosen or obtained in dimensioning the circuit parameters are reported in Table ST1.

| **Compartment #** | **1** | **2** | **3** | **4** |
|---|---|---|---|---|
| $\mathbf{g_{eq}}$ [nS] | 72.5 | 41.2 | 23.4 | 13.3 |

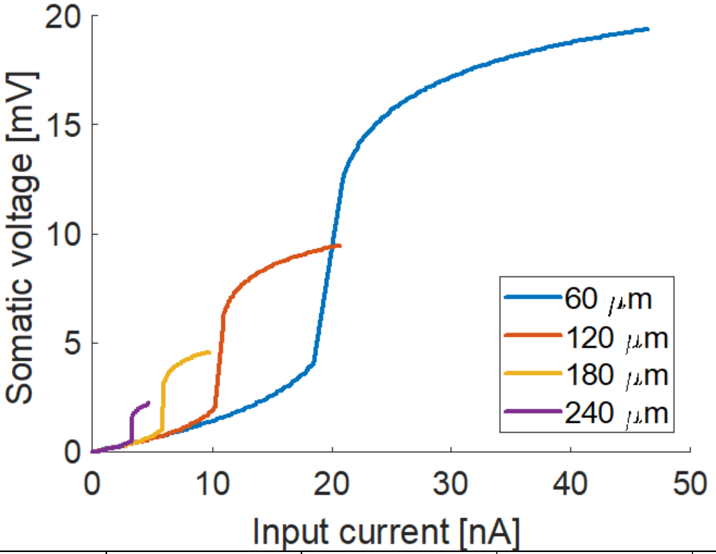


| | | | | |
|---|---|---|---|---|
| $\gamma_{NMDA,n}$ [nS/a.i.u.] | 173 | 86.8 | 43.6 | 21.9 |
| $I_{ex\text{-}th,n}$ [a.i.u.] | 1.49 | 1.67 | 1.88 | 2.11 |
| $I_{th,n}$ [nA.] | 2 | 1.2 | 0.74 | 0.45 |
| $g_L$ [nS] | 10.2 | 4.3 | 2.5 | 2.8 |
| $g_a$ [nS] | 81.5 | 65.2 | 37.1 | 21.1 |
| $g_{SOMA}$ [nS] | 165.7 | | | |
| eff. $V_{peak,n}$ [mV] | 19.4 | 9.5 | 4.6 | 2.2 |

**Table ST1** – Circuit parameters (both selected and calculated) vs compartment number used in the simulations.

Figure SF-4 shows the somatic responses to excitatory currents at the four compartments as from the calculations above.

**Figure SF-4** – Somatic voltage obtained upon application of excitatory inputs at a single location, vs the distance of the location from the soma. As expected, the peak amplitude increases (and the threshold decreases) vs the distance.

*Equalization of cell responses*

Since the cells have variable excitatory/inhibitory input balance, the somatic voltage range varies considerably – in natural neurons other equalizing mechanisms may be present including, possibly, lateral connections, which are not accounted for, here.

To equalize the responses of the cells, the 80th percentile of the responses to the grating stimuli was determined for each cell (Figure SF-5): The relationship with $\sigma_{rel}$ is evident – lower- $\sigma_{rel}$ cells, with less effective inhibition, have larger somatic response on average. A multiplying constant was then applied to the somatic voltage to set the 80th-percentile response to the threshold (15 mV above rest, or -55 mV). The correction was applied to all the figures in the main text.

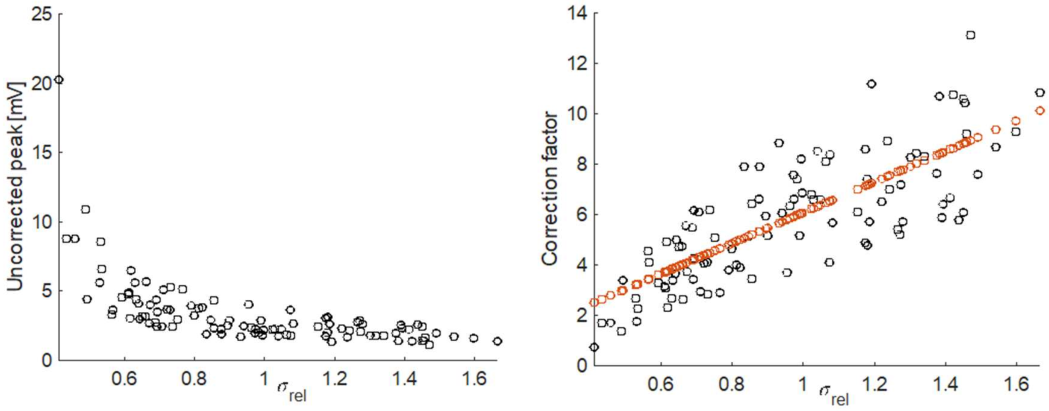


**Figure SF-5** – (left) 80th percentile of the uncorrected cell responses to grating stimuli vs $\sigma_{rel}$. (right) This was linearly corrected to be the firing threshold values (-55 mV) (black: punctual corrections; red: interpolated corrections applied to cells)

#### *SI-2 Angular distributions of excitatory and inhibitory inputs*

The distribution parameters ($\sigma_{in}$ in particular) were determined basing on the cell's radius, $\sigma_{ex}$ (40 to 50 degrees) and the excitatory-to-inhibitory peak ratio (0.3 to 0.7). Once fixed the total number of connections (300), the number of excitatory connections required if the radius were minimum, $r_0$, was calculated. Let

$$p_{ex}^{0} = c_{ex/in} e^{-\frac{\vartheta^2}{2\sigma_{ex}^2}} \text{ and } p_{in}^{0} = e^{-\frac{\vartheta^2}{2\sigma_{in,max}^2}} \quad \text{(SE22)}$$

be the two corresponding, non-normalized Gaussian distributions centered around a preferred angular orientation (θ=0, with no loss of generality). The second has maximum standard deviation (70 degrees) as it would be at the minimum radius. In this case, the number of excitatory connections would be obtained integrating the two distributions, and considering that

$$\frac{n_{in}^0}{n_{ex}^0} = \frac{\sigma_{in,max}}{c_{ex/in}\sigma_{ex}}, \quad \text{(SE23)}$$

so that

$$n_{ex}^0 = \frac{1}{1+\frac{n_{in}^0}{n_{ex}^0}} n_{tot}, \quad \text{(SE24)}$$

where $n_{tot}$=300 is the total number of connections per cell.

The radius was chosen uniformly between $r_0$ and (about) $r_0\sqrt{2}$ – that is, between 11 and 16 pixels – and the effective number of excitatory and inhibitory connections was calculated, the first assuming uniform density and the second by difference with respect to the total:

$$n_{ex} = n_{ex}^0 \left(\frac{r}{r_0}\right)^2 \text{ and } n_{in} = n_{tot} - n_{ex}. \quad \text{(SE25)}$$

Finally, the standard deviation of the inhibitory distribution was obtained readapting (SE18):

$$\sigma_{in} = c_{ex/in} \frac{\sigma_{in} n_{in}}{n_{ex}}. \quad \text{(SE26)}$$

*SI-3 Gabor filters.*

Three examples of Gabor filters, built as in Methods, are sketched in Figure SF-6, for different orientations and polarities:

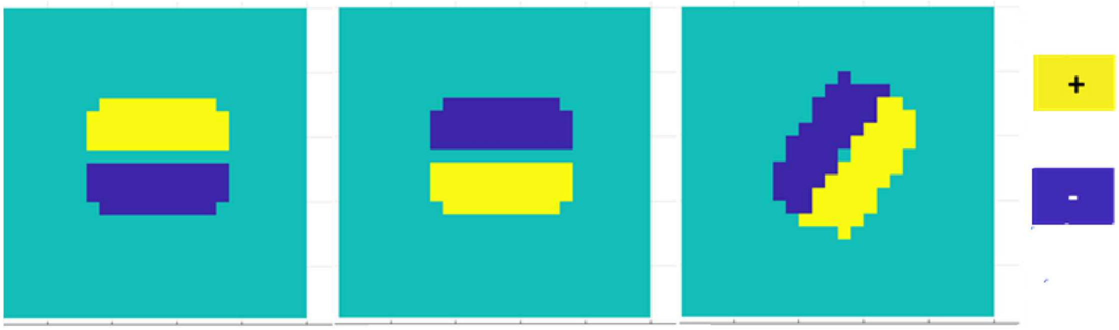


**Figure SF-6** – Three of the 20 Gabor filters used, for 0 degrees (left and middle) and 54 degrees (right) respectively.

**Figure SF-7** (cont'd) – Cells 1 to 12

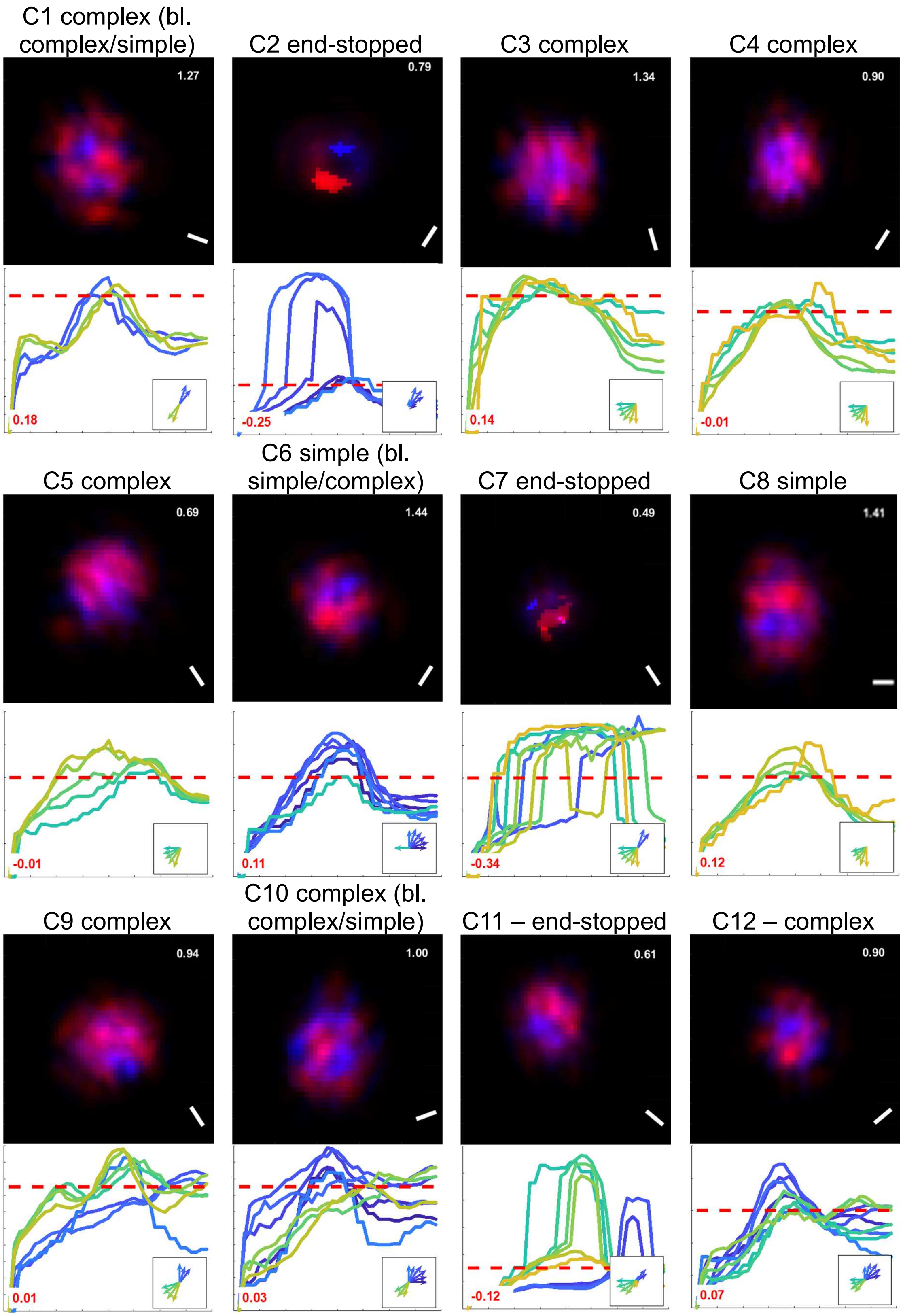

**Figure SF-7** (cont'd) – Cells 13 to 24

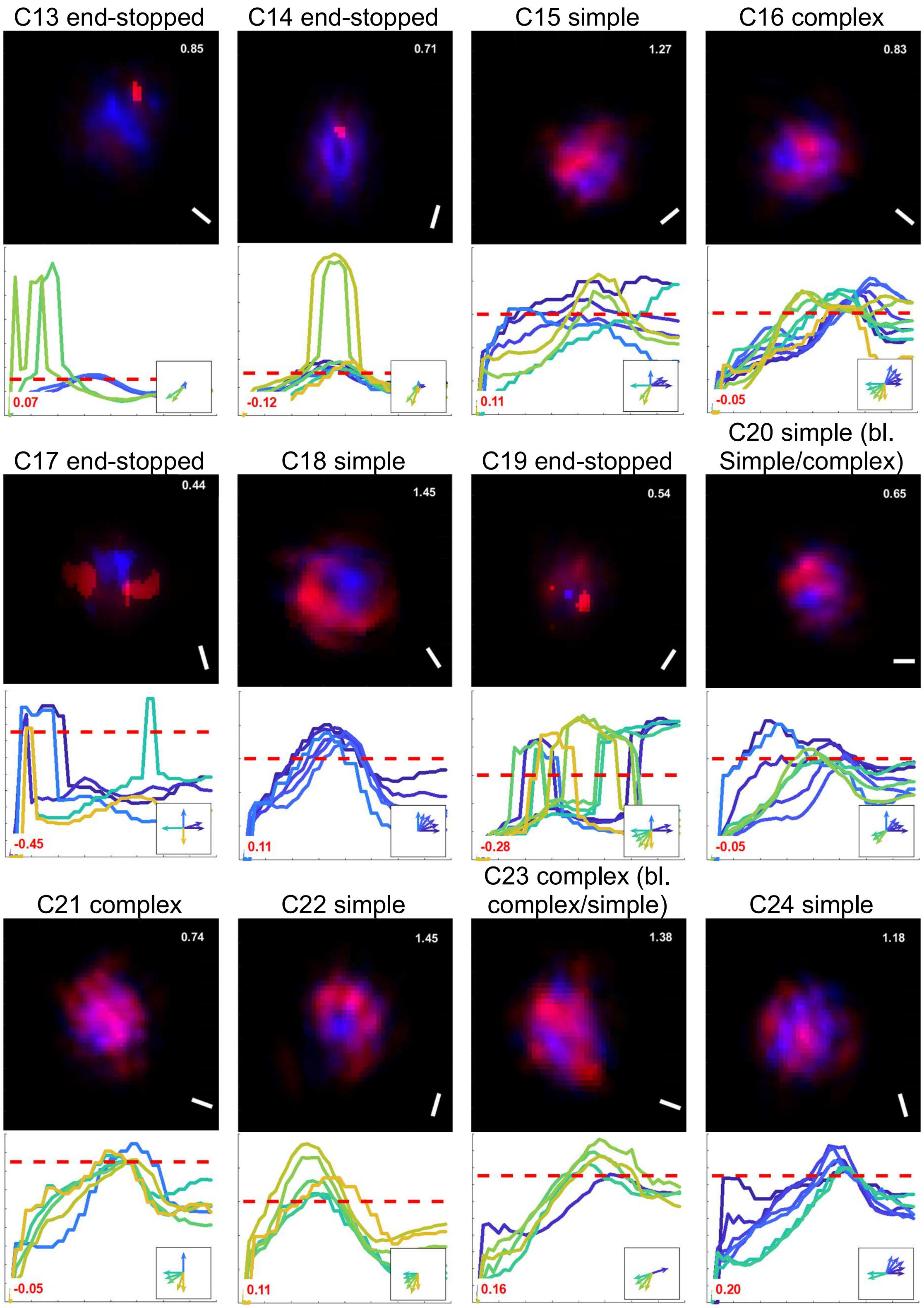

**Figure SF-7** (cont'd) – Cells 25 to 36

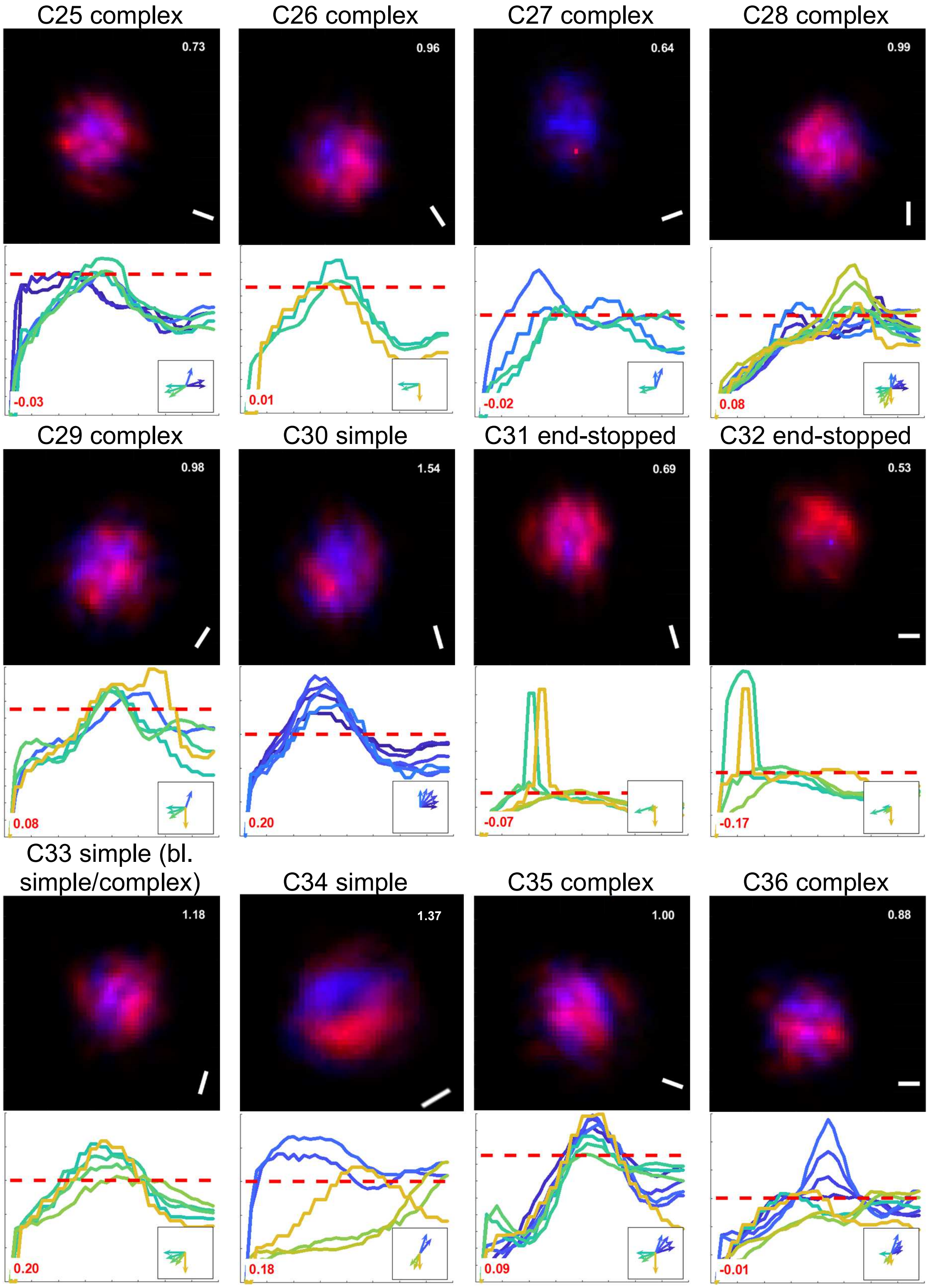

**Figure SF-7** (cont'd) – Cells 37 to 48

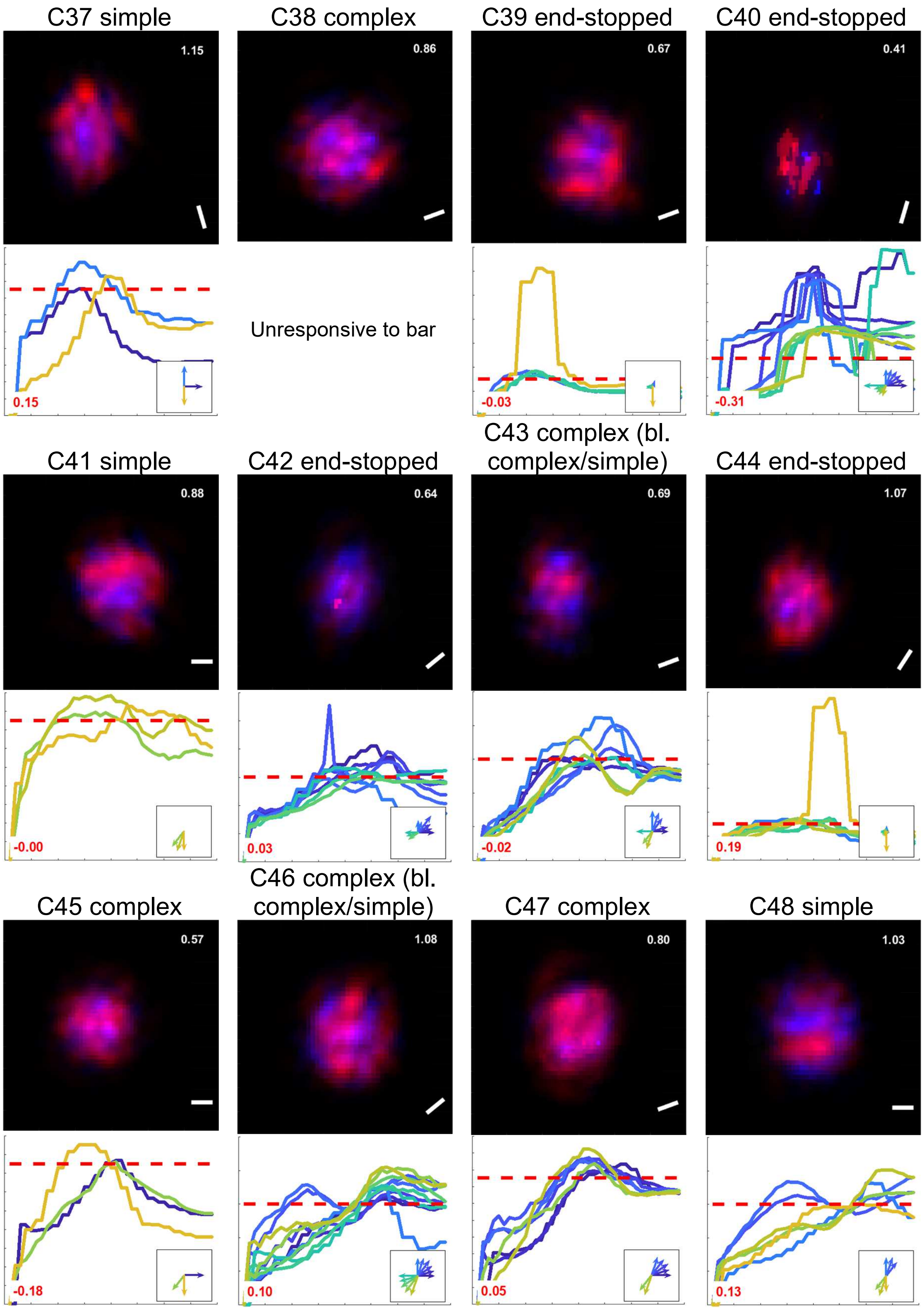

**Figure SF-7** (cont'd) – Cells 49 to 60

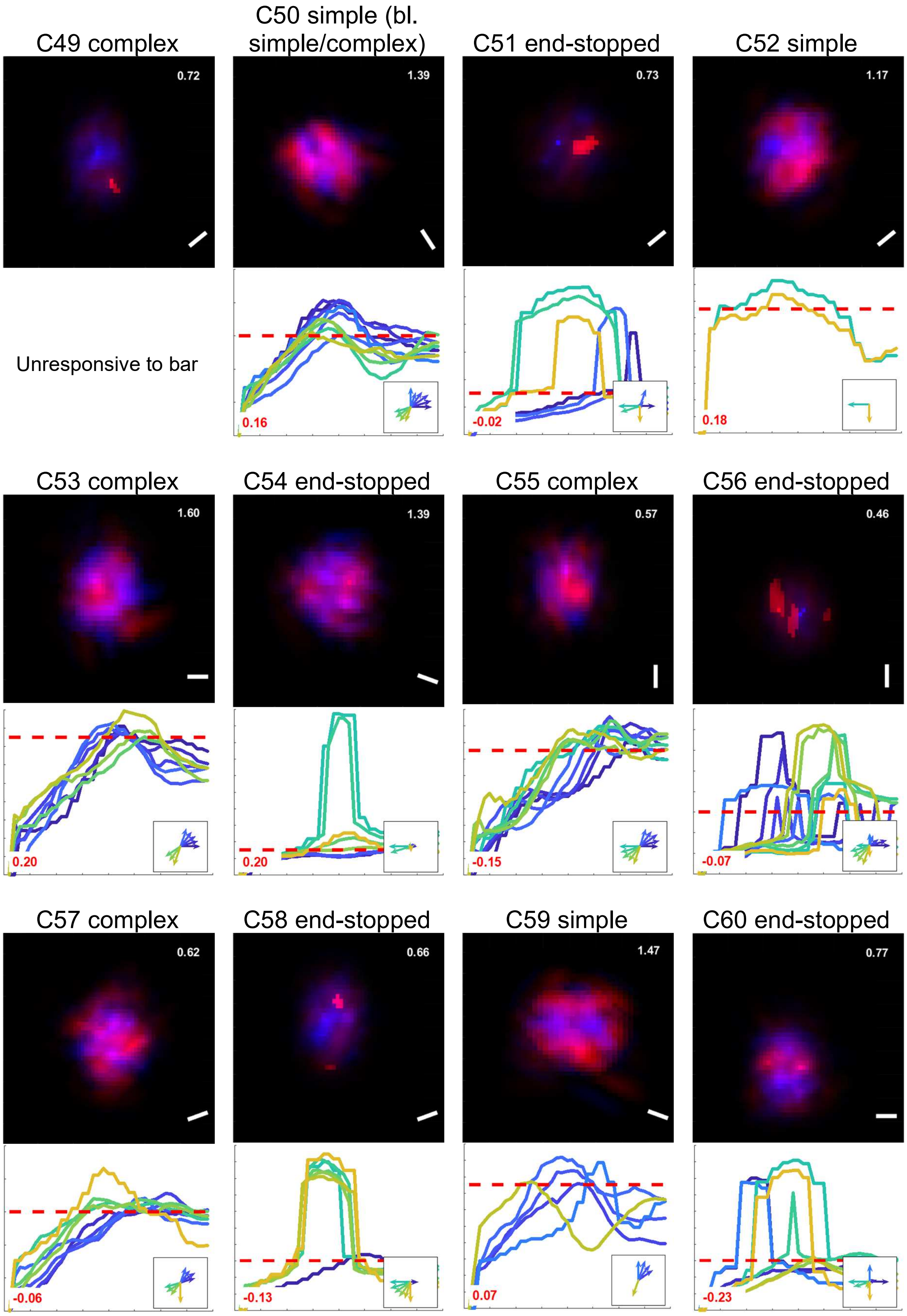

**Figure SF-7** (cont'd) – Cells 61 to 72

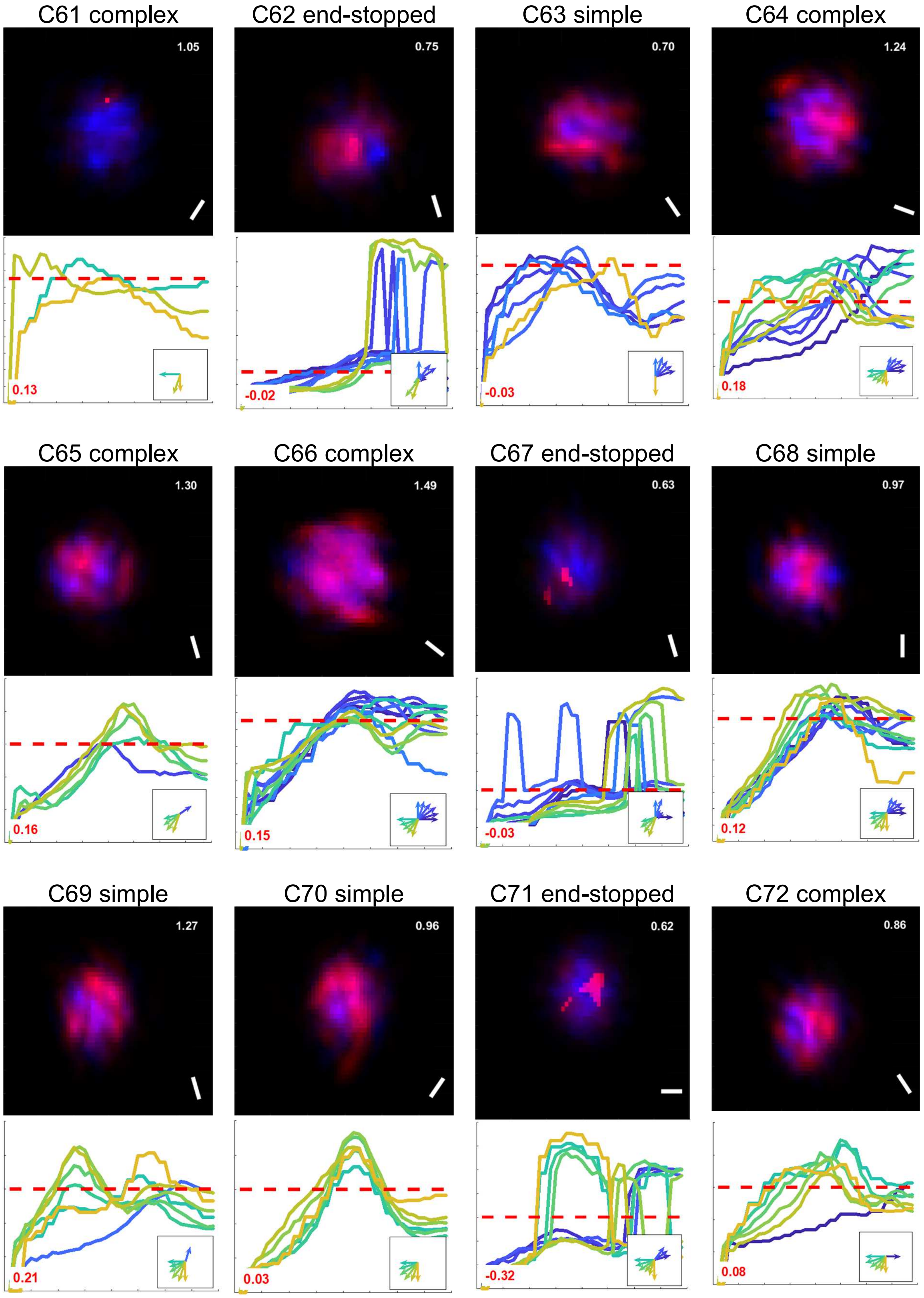

**Figure SF-7** (cont'd) – Cells 73 to 84

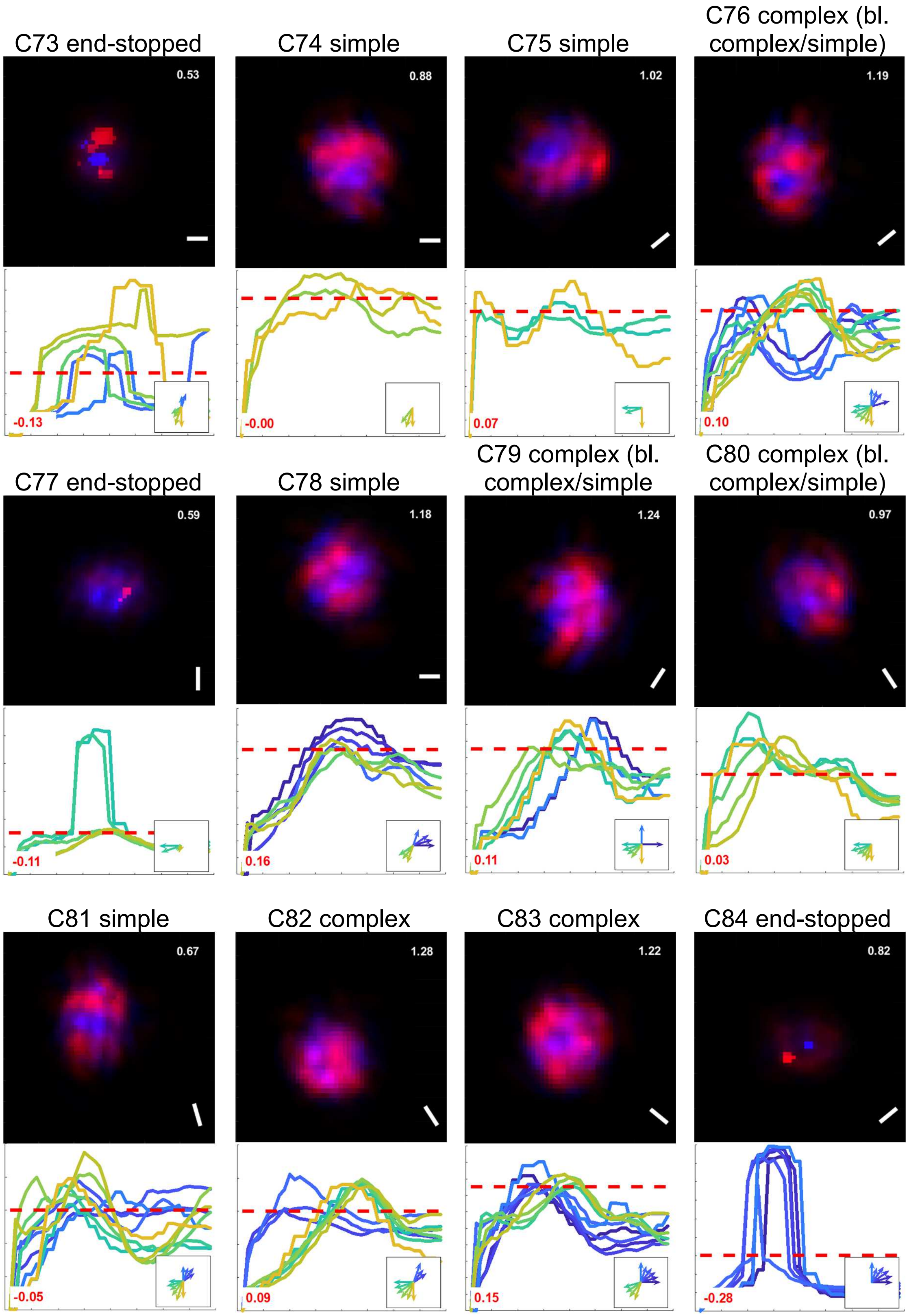

**Figure SF-7** (cont'd) – Cells 85 to 96

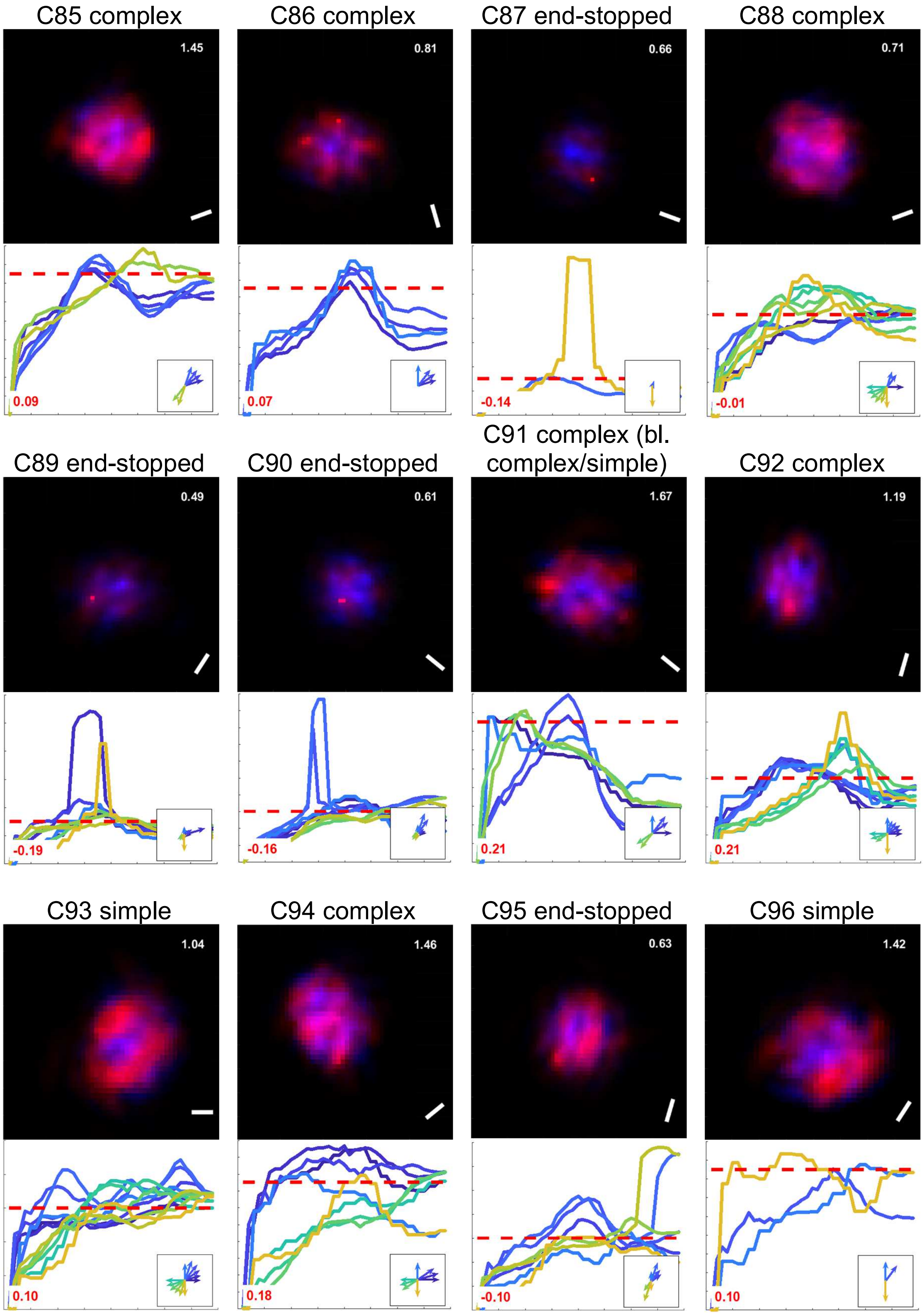

**Figure SF-7** – Cells 97 to 100

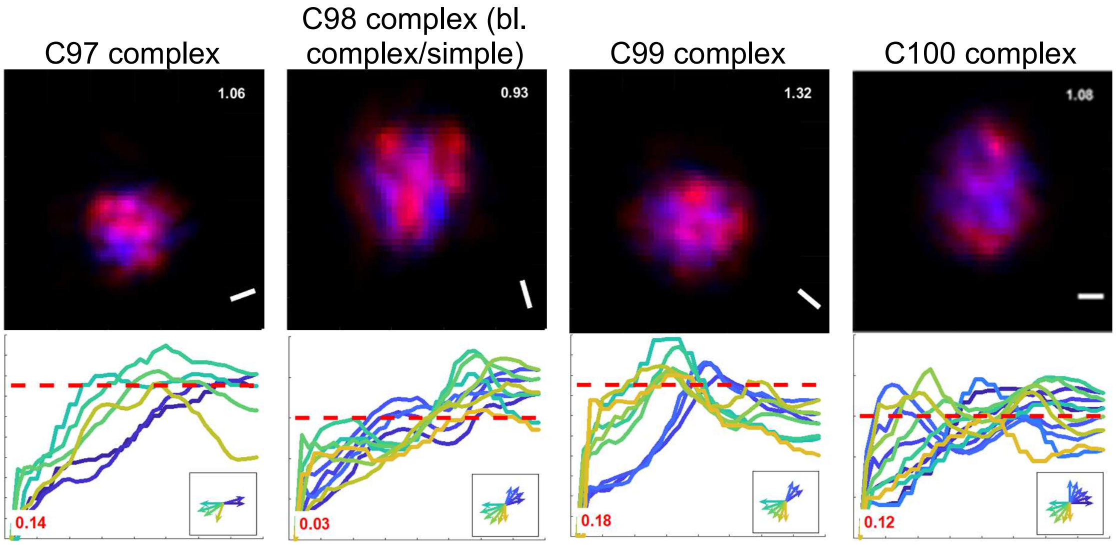


**Figure SF-7 –** For each cell, the spot map (top) and responses to bar stimuli of different orientations and lengths (bottom) are shown. In the spot map, $\sigma_{rel}$ is reported in the upper right corner and the assigned preferred orientation (from the input distributions) is indicated by a bar in the lower right corner. In the bar-response plots, only responses exceeding the −55 mV threshold are shown (red dotted line). The red number in the lower left corner of each panel reports the corresponding effective membrane orientation selectivity index, $mOSI_{eff}$. In the lower right inset, the directions of bars capable of eliciting a response are represented as arrows whose length is proportional to the maximum response. Each panel also reports the cell identifier and its classification as simple, complex, or end-stopped (for example, “C18 simple”, “C47 complex”, “C2 end-stopped”). For a few neurons, the classification is additionally marked as borderline, using the notation “(bl. cat1/cat2)” to indicate behavior intermediate between categories (for example, “C50 simple (bl. simple/complex)”). These borderline cases illustrate the gradual transitions between response types in the model and the qualitative nature of the classification criteria.

**Figure SF-8** – Somatic voltage of three cells as a function of grating stimuli presented at different orientations (arranged by columns), and with different periods and phases (as indicated in the upper left picture).

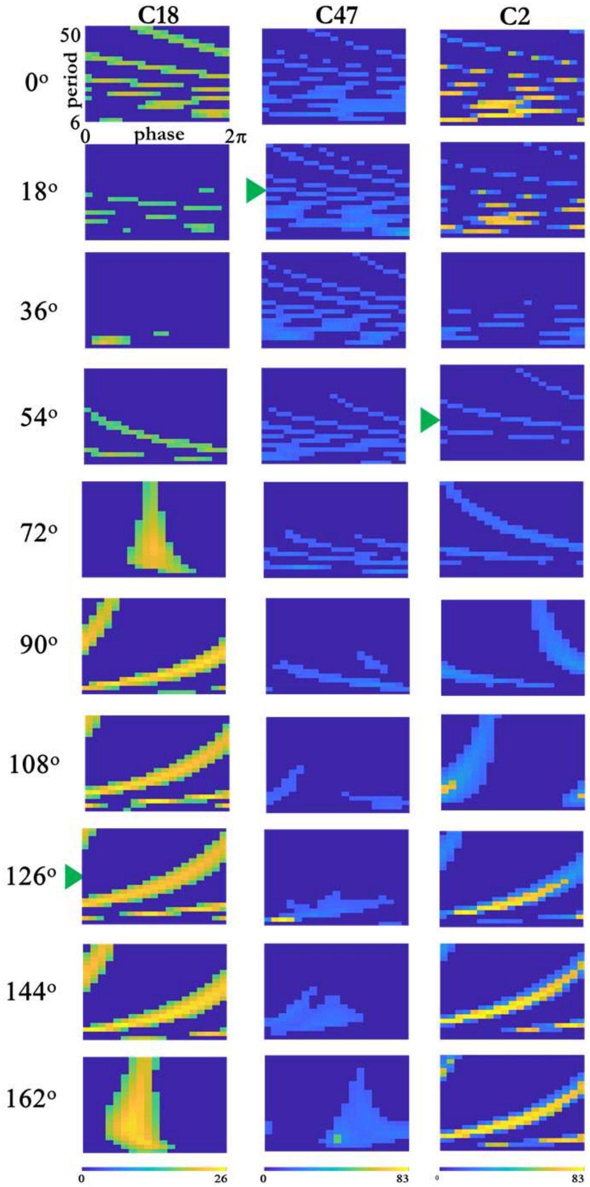

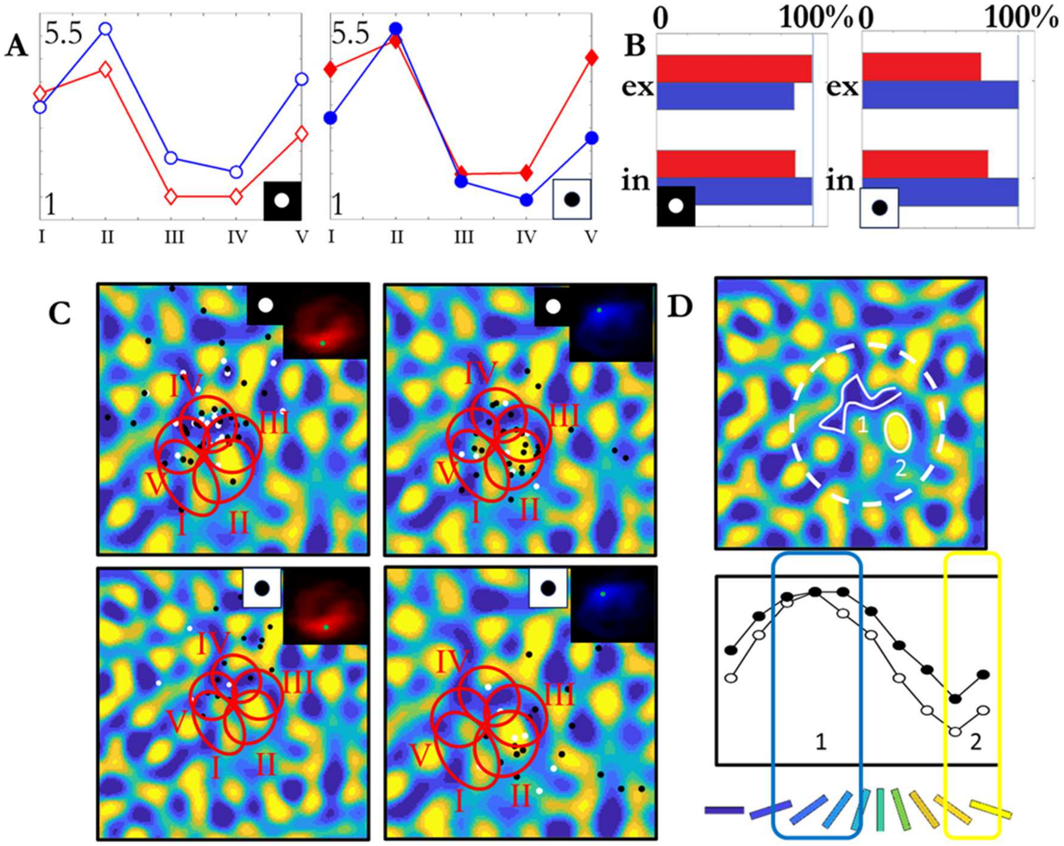


**Figure SF-9** – Analysis of simple cell C34, carried out as for C18 (coding as in Figure 4, main text). **A.** The inhibitory-to-excitatory ratios are close to those of C18 producing the same differentiation between ON and OFF regions. **B** Besides the increase in inhibitory input from ON to OFF for both stimuli, in this case there is also some decrease in excitation for the bright spot. **C** and **D** At the peak of the ON region, the bright spot stimulates many connections, a large fraction of which is excitatory (area 1, dense and centered around the preferred stimulation), mostly in the proximal compartment of dendrite IV. Conversely, the dark spot, at its maximum effectiveness, stimulates connections mostly in the proximal compartment of dendrite II, corresponding to region 2, where inhibition dominates but is collected less effectively by the dark Gabor ring.

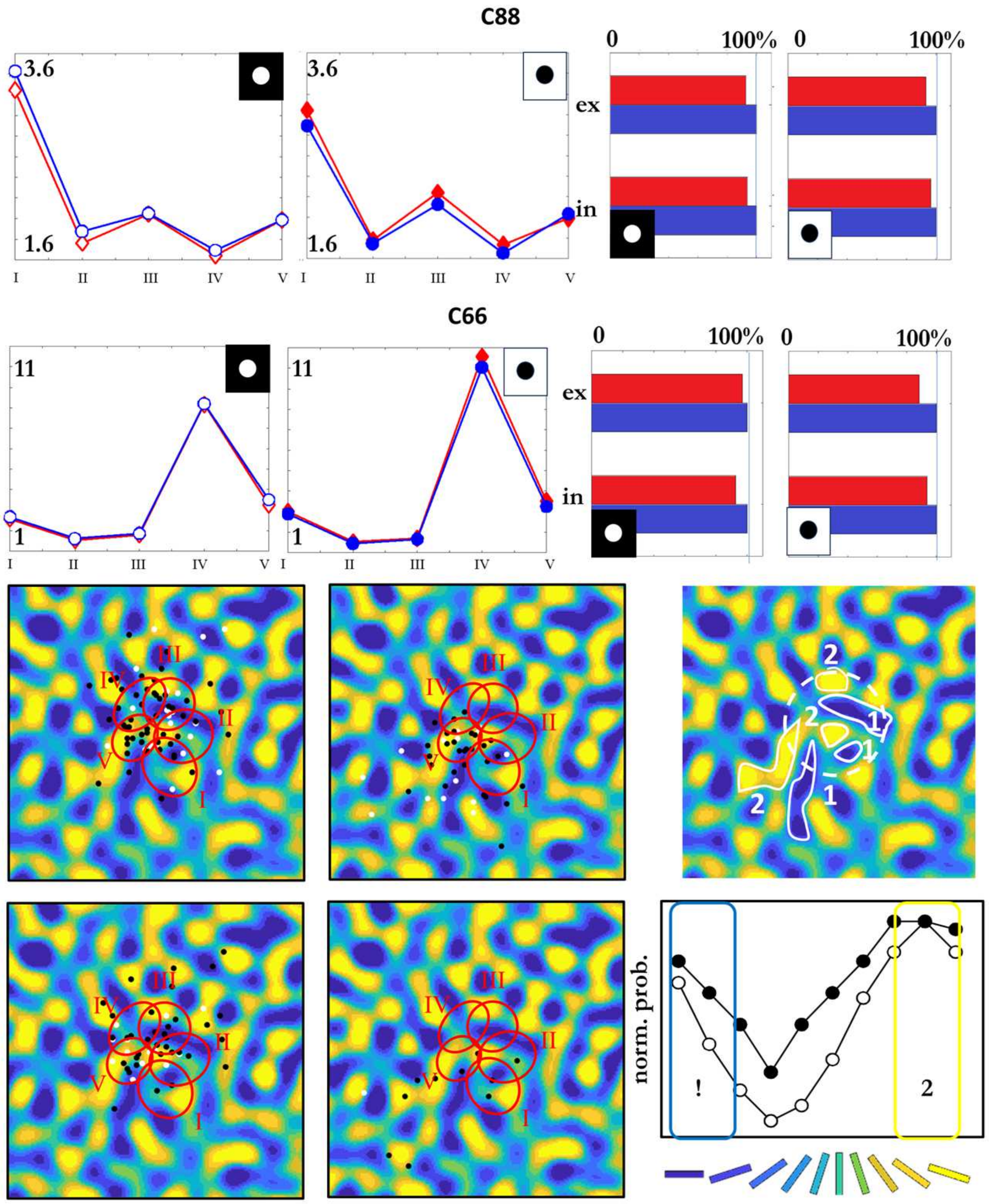


**Figure SF-10** – Analysis of complex cells C66 and C88. C88 behaves as a complex-like cell for the same reason as C47, namely a small $\sigma_{rel}$. Conversely, C66 is complex-like despite a large $\sigma_{rel}$≈1.49. In this case, the two central orientation subregions 1 and 2 (shades of yellow and blue, respectively) are arranged in bands whose spatial periodicity is smaller than the size of the dendritic compartments. As a result, individual compartments often receive input from both subregions, and there are no significant differences in connection density that would separate ON and OFF regions, leading to substantial ON/OFF overlap. This example confirms that the detailed local structure of the orientation map, and the resulting dendritic activation patterns, can override the influence of global input statistics as summarized by $\sigma_{rel}$.
